# Population genetic analysis of *Ophidiomyces ophidiicola*, the causative agent of snake fungal disease, indicates recent introductions to the USA

**DOI:** 10.1101/2022.03.23.485546

**Authors:** Jason T. Ladner, Jonathan M. Palmer, Cassandra L. Ettinger, Jason E. Stajich, Terence M. Farrell, Brad M. Glorioso, Becki Lawson, Steven J. Price, Anne G. Stengle, Daniel A. Grear, Jeffrey M. Lorch

## Abstract

Snake fungal disease (SFD; ophidiomycosis), caused by the pathogen *Ophidiomyces ophidiicola* (*Oo*), has been documented in wild snakes in North America and Eurasia, and is considered an emerging disease in the eastern USA. However, a lack of historical disease data has made it challenging to determine whether *Oo* is a recent arrival to the USA or whether SFD emergence is due to other factors. Here, we examined the genomes of 82 *Oo* strains to determine the pathogen’s history in the eastern USA. *Oo* strains from the USA formed a clade (Clade II) distinct from European strains (Clade I), and molecular dating indicated that these clades diverged too recently (∼2,000 years ago) for transcontinental dispersal of *Oo* to have occurred via natural snake movements across Beringia. A lack of nonrecombinant intermediates between clonal lineages in Clade II indicates that *Oo* has actually been introduced multiple times to North America from an unsampled source population, and molecular dating indicates that several of these introductions occurred within the last few hundred years. Molecular dating also indicated that the most common Clade II clonal lineages have expanded recently in the USA, with time of most recent common ancestor mean estimates ranging from 1985-2007 CE. The presence of Clade II in captive snakes worldwide demonstrates a potential mechanism of introduction and highlights that additional incursions are likely unless action is taken to reduce the risk of pathogen translocation and spillover into wild snake populations.

## INTRODUCTION

Emerging infectious diseases (EIDs) are increasingly recognized as having major impacts on wildlife conservation and global biodiversity [1–3]. Anthropogenic activities resulting in movement of pathogens to new locations and environmental change are the driving forces behind the emergence of many wildlife diseases [1,2,4–6]. Due to several characteristics that make them pernicious pathogens (e.g., high virulence, ability to establish environmental reservoirs, and broad host range), fungi have been responsible for some of the most consequential of these EIDs [7]. For example, the emergence of white-nose syndrome, a disease of hibernating bats caused by *Pseudogymnoascus destructans*, has caused population declines in excess of 90% for several bat species across eastern North America [8]. Similarly, *Batrachochytrium dendrobatidis*, the causative agent of chytridiomycosis, has been linked to the global decline of hundreds of amphibian species and the extinction of at least 90 species [9].

Mechanisms for disease emergence typically fall into one of two broad categories: (1) introduction of an exotic pathogen into a naïve host population (“novel pathogen hypothesis”) or (2) *in situ* emergence of a native pathogen due to changes in environmental, host, or pathogen characteristics that alter disease ecology (“endemic pathogen hypothesis”) [10]. Understanding mechanisms of disease emergence is key for assessing the threat of a disease to host populations and predicting pathogen spread, as well as for devising disease management strategies. For example, while targeted containment or eradication efforts may be effective against novel pathogens, successful management of endemic pathogens is more likely to involve manipulation of factors (e.g., environmental, host) that contribute to disease outbreaks [10].

Despite the importance of identifying the reason for emergence of a disease, the origin of fungal pathogens is often elusive, and the sources of many important fungal (and fungal-like) pathogens remain unknown (e.g., *Bretziella fagacearum*, causative agent of oak wilt [11]; *Ophiognomonia clavigignenti-juglandacearum*, causative agent of butternut canker [12]; *Phytophthora ramorum*, causative agent of sudden oak death [13]). A lack of historical samples and surveillance, temporal delays between actual emergence and recognition of the disease, and the potential for multiple introductions and recombination between strains can make it difficult to ascertain the extent of a fungus’ geographic distribution and whether it is native or introduced to a region [14–16].

Snake fungal disease (SFD or ophidiomycosis) is often cited as a fungal EID afflicting wildlife [17, 18], although the extent to which SFD is actively emerging is difficult to quantitatively assess [19, 20]. The disease first gained attention in 2008 when severe fungal infections manifested in a well-studied population of eastern massasauga rattlesnakes (*Sistrurus catenatus*) in Illinois, USA [21]. Subsequent investigations revealed that SFD was widely distributed in the eastern USA and Great Lakes region of Canada [18,20,22]. The fungus *Ophidiomyces ophiodiicola* (*Oo*) is the causative agent of SFD [23, 24]. Although *Oo* is not known to infect other animals, the fungus has a broad host range among snakes and most snake taxa are predicted to be susceptible [25]. However, disease severity and outcomes of infection are variable [18], and the broader conservation impacts of SFD are difficult to predict, in part because the extent of, and mechanism behind, the disease’s emergence within snake populations is unknown.

SFD is now endemic in wild snakes in eastern North America [18, 20]. However, existing evidence as to the origin of *Oo* in the eastern USA is ambiguous. Under the novel pathogen hypothesis, we would predict that *Oo* arrived in the USA relatively recently, perhaps in the order of decades prior to the increase in cases of SFD reported during the early 2000s [18, 21]. Indeed, the detection of genetically unique and diverse strains of *Oo* from wild snakes in Eurasia [26, 27] is consistent with this hypothesis, as it indicates that *Oo* may have originated from outside North America. In contrast, the endemic pathogen hypothesis would be consistent with a much older arrival of *Oo* in the USA, with the recent increase in SFD cases linked to greater awareness or environmental changes rather than a new introduction. *Oo* is thought to be a specialized pathogen of snakes with limited ability to survive as an environmental saprobe [28]. Thus, under the endemic pathogen hypothesis *Oo* most likely evolved in North America or arrived in the USA through the natural migration of snakes from Eurasia to North America via Beringia, which occurred in the span of 27-55 million years ago [29, 30]. Detections of SFD in snakes in the USA as early as 1945 [31] and the lack of a documented wave-like spread pattern after the reported emergence of the disease in the early 2000s [18] could support the endemic pathogen hypothesis. However, a lack of sufficient historical data on the disease has made it difficult to trace the origins of *Oo* in North America.

Phylogenetic and population genetic analyses based on whole genome sequence data provide powerful tools for inferring the origins of fungal pathogens in the absence of historical samples [32]. For example, Drees et al. [33] demonstrated that isolates of *P. destructans* from North America exhibited minimal genetic diversity, consistent with clonal expansion following a single introduction event from Eurasia. In the case of *B. dendrobatidis*, analyses based on whole genome sequence data ultimately revealed southeast Asia as the likely source of the fungus - a finding that was long evasive due to multiple introduction events involving several genetic lineages and recombination of those lineages upon recontact [15, 16]. Here we report full genome sequences of 82 strains of *Oo* and phylogenetic and population genetic analyses that explore the origins of *Oo* in North America. Our findings indicate that strains of *Oo* in the eastern USA are primarily represented by four clonally-expanded lineages or hybrids between those lineages, and that the ancestors of these clonal lineages arrived in the region relatively recently.

## RESULTS

### Whole genome sequencing and annotation

Using the Illumina platform, we obtained whole genome shotgun sequencing data for 82 strains of *Oo*, with 1.2–10.8 M paired end reads obtained for each (median = 2.4 M). After quality filtering and trimming, this provided 9.3-202.4x average genome coverage depth (median = 23.5x; Table S1). Using these data, we assembled both the nuclear and mitochondrial genomes *de novo*, with 81–1,350 nuclear contigs per strain (median = 411) and a single mitochondrial contig per strain (length = 42,585 - 54,364). We used the *de novo* contigs to predict open reading frames and protein sequences (6,901–7,190 predicted open reading frames per strain).

### Mating type

Predicted mating type proteins identified in *Oo* shared 43.6–61.7% amino acid identity with the conserved portion of the alpha box mating type protein MAT1-1 and 44.4–62.7% amino acid identity with the conserved portion of the high mobility group (HMG) domain-containing MAT1-2 protein of other Onygenalean fungi. Each *Oo* strain possessed either a putative *MAT1-1* or *MAT1-2* locus, consistent with a heterothallic mating system. Both mating type idiomorphs were observed in isolates from wild snakes in the USA and in captive snakes (Table S2). However, all four European isolates sampled possessed the *MAT1-2* idiomorph. To further explore mating type idiomorphs present in Europe, we screened nine additional strains of *Oo* isolated from wild snakes in the United Kingdom that were not included in the whole-genome sequence analysis for *MAT1-1* and *MAT1-2* using targeted PCR assays (see Supporting Text). Only *MAT1-2* was detected in these nine strains.

### Phylogenetic Analysis

To obtain a genome-wide view of the evolutionary relationships among the 82 *Oo* strains, we generated a maximum-likelihood phylogeny using a concatenated amino acid alignment of 5,811 proteins (Fig. S1). This represented ∼81-84% of the annotated protein coding genes for each strain, and the overall topology, which consisted of three, well-supported main clades, was consistent with previously published phylogenies for *Oo* [26, 27]. One of these clades (Clade I) contained all four strains isolated from wild snakes in Europe; a second clade (Clade II) contained all 65 strains isolated from wild snakes in North America as well as 10 strains from captive snakes; and the third clade (Clade III) included three strains, all collected from snakes in captivity. Notably, strains from Clades II and III were observed in captive snakes from all three sampled continents (North America, Europe, and Australia). Consistent with previous studies, we found that Clade III was an outgroup to Clades I and II; however, our full genome analysis indicated a much higher level of genetic divergence than previously reported. The two protein coding genes used in previous phylogenetic analyses of *Oo* (actin and translation elongation factor 2L [26, 27]) exhibited average levels of divergence between Clades III and I/II (0.016% and 0% at the amino acid level) that were lower than ∼97-98% of the orthologs we examined (Fig. S1B). Average per gene amino acid divergence between Clades III and I/II was 2.6% (median = 1.9%).

We also observed the same three well-supported clades in a phylogeny based on a nucleotide-level single nucleotide polymorphism (SNP) alignment across the full mitochondrial genome (Fig. S1C). However, while mitochondrial genetic diversity within each clade was generally low (<0.028%, <0.018%, and ≤0.133% divergence for Clades I–III, respectively), one strain from Clade II (NWHC 44736-75) exhibited a high level of genetic divergence compared to the other members of this clade (0.20–0.21%). Levels of genetic divergence between strain NWHC 44736-75 and Clades I and III were even higher (0.32–0.35% and 1.53–1.56%, respectively). Combined with phylogenetic placement based on the nuclear genome, this is consistent with strain NWHC 44736-75 being of hybrid origin between Clade II and a lineage that is not represented in our dataset.

To better resolve relationships within the North American and European clades (i.e., through inclusion of more informative sites), we removed the three highly divergent strains from Clade III and performed a nuclear genome-wide phylogenetic analysis at the nucleotide-level. This phylogeny was based on a 19,699,744 base pair (bp) alignment, which included 189,199 variable sites. For Clade I, which includes the four European strains, we observed strong concordance between the patterns of genetic divergence in the nuclear and mitochondrial genomes (Fig. 1). However, no such concordance was observed within Clade II, which includes all of the North American strains.

**Fig. 1.**
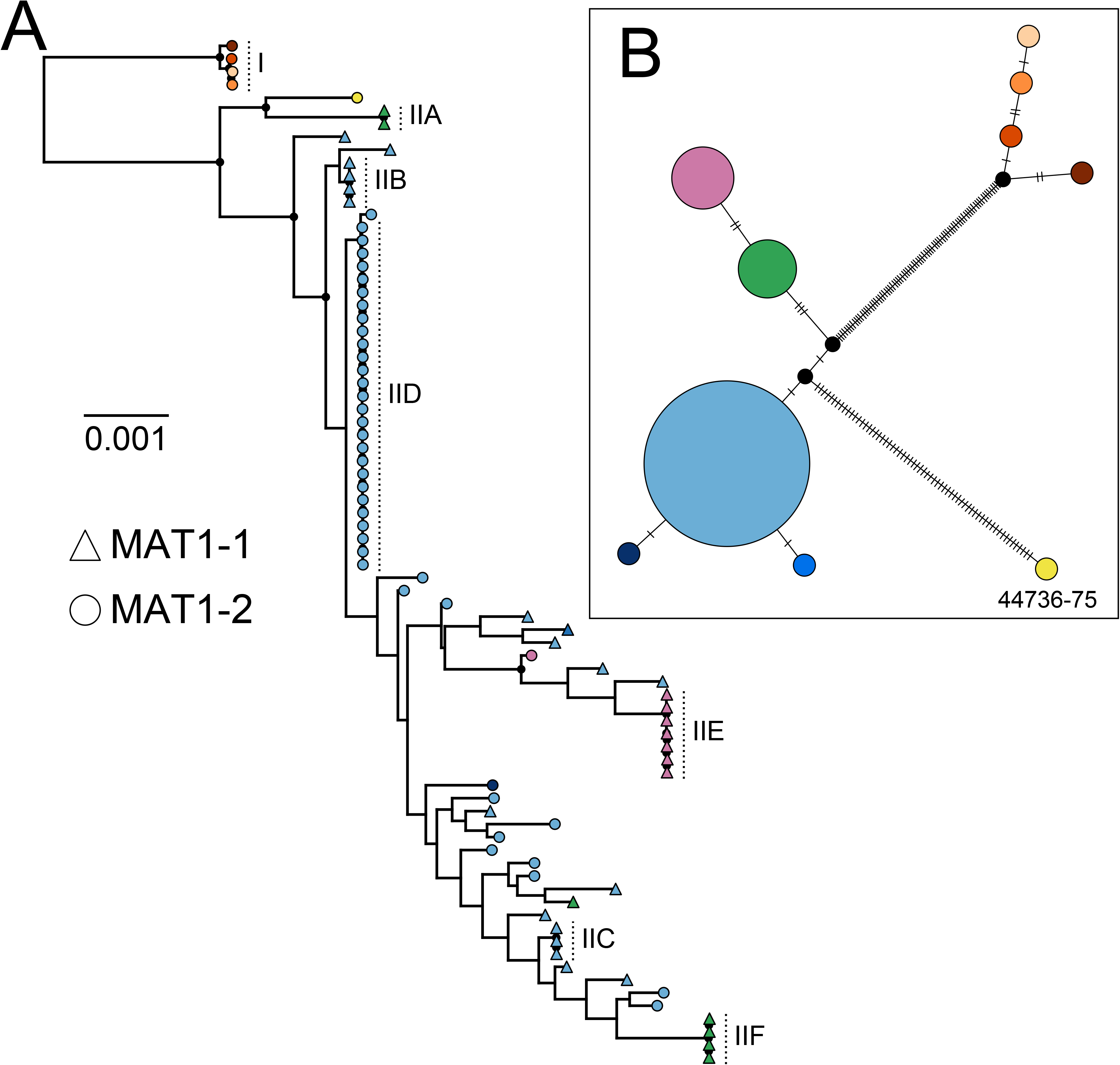
All North American isolates of *Ophidiomyces ophidiicola* form a well-supported clade, but there is discordance between genetic relationships within the nuclear (A) and mitochondrial (B) genomes. (A) Maximum-likelihood phylogeny including all 79 strains from Clades I and II and based on an alignment of 19,699,744 base pairs, including 189,199 single nucleotide polymorphisms (SNPs). Tip shapes indicate the mating type of each strain and colors correspond to the mitochondrial genotypes presented in (B). Filled black circles indicate nodes with bootstrap support ≥ 90. Vertical dotted lines and labels indicate the strains of Clade I, as well as the putative clonal lineages within Clade II. (B) A median-joining haplotype network including all 79 strains from Clades I and II and built from 159 SNPs present in the mitochondrial genome (“best” SNPs from NASP). Haplotype colors correspond to the tip colors from (A). Each tick mark corresponds to a single SNP difference and black circles represent inferred, but unsampled nodes.

While we observed several apparently clonal lineages within Clade II, many of the strains from this clade were highly divergent from all other strains. In contrast, just three mitochondrial genotypes accounted for 96% of all Clade II strains (72/75), all three of these genotypes were observed in strains that exhibited substantial levels of genetic divergence within the nuclear genome, and all three exhibited polyphyletic nuclear relationships (Fig. 1). Additionally, despite long branch lengths in the nuclear phylogeny, most of the nodes within Clade II exhibited low bootstrap support. These patterns all indicated a history of recombination within Clade II (see below). However, we did not find any evidence of recombination among strains contained within the same putatively clonal lineage. Specifically, the nuclear phylogeny supported six apparently clonal lineages (Fig. 1, IIA-IIF), each of which included 2-27 *Oo* strains. All of the strains within a given clonal lineage shared an identical mitochondrial genotype and had the same mating type locus. Additionally, we found little, if any, evidence for homoplasy within these lineages based on a parsimony analysis with PAUP* (consistency indexes = 0.93–1, only run for the four clonal lineages that contained ≥4 strains) (Table S3). We also saw little evidence of homoplasy among the strains of Clade I (consistency index = 0.99).

### Recombination Analyses

We observed a very low consistency index (0.25) in our analysis that included all 75 *Oo* strains from Clade II (Table S3), which is indicative of a history of recombination. Specifically, this indicated that the most parsimonious tree for this dataset required four times the minimum number of required changes because of the presence of homoplasy. For each of the 17 scaffolds with >50,000 high confidence base calls, we also broadly assessed recombination using the Phi test [34]. We detected widespread evidence of recombination among the *Oo* strains of Clade II, with significant evidence for recombination observed on each of the 17 scaffolds, even after correcting for multiple tests (Fig. S2).

To further explore the genetic relationships among *Oo* strains and to look for evidence of recent recombination within Clade II, we generated a co-ancestry matrix using ChromoPainter [35]. A principal component analysis (PCA) based on our co-ancestry matrix revealed patterns consistent with recent recombination among three of the clonal lineages in Clade II: IID, IIE, and IIF (Fig. 2B). Specifically, we found that 25 of the *Oo* strains fell along a gradient between IID and IIF, while eight strains fell along a gradient between IID and IIE, and one additional strain (NWHC 24411-01) appeared to fall along one of these gradients, but very close to the IID clonal lineage. These patterns are consistent with recent hybridization between these lineages as all IID strains possessed the *MAT1-2* mating type locus, whereas the strains from IIE and IIF all possessed the *MAT1-1* mating type locus. The putative hybrids contained a mixture of these two mating types. Only two examples of clonal expansion within a recombinant lineage were observed (IIB and IIC). In both instances, the clonal strains within the recombinant lineages were collected from the same location (Norfolk County, Massachusetts, USA for IIB; Virginia Beach, Virginia, USA for IIC), consistent with limited geographic distributions; however, even with modest sample sizes (3-4 strains), each of these clonal lineages was isolated from multiple snake species. All other hybrids were represented by a single isolate and appeared to indicate separate recombination events (Fig. 2C).

**Fig. 2.**
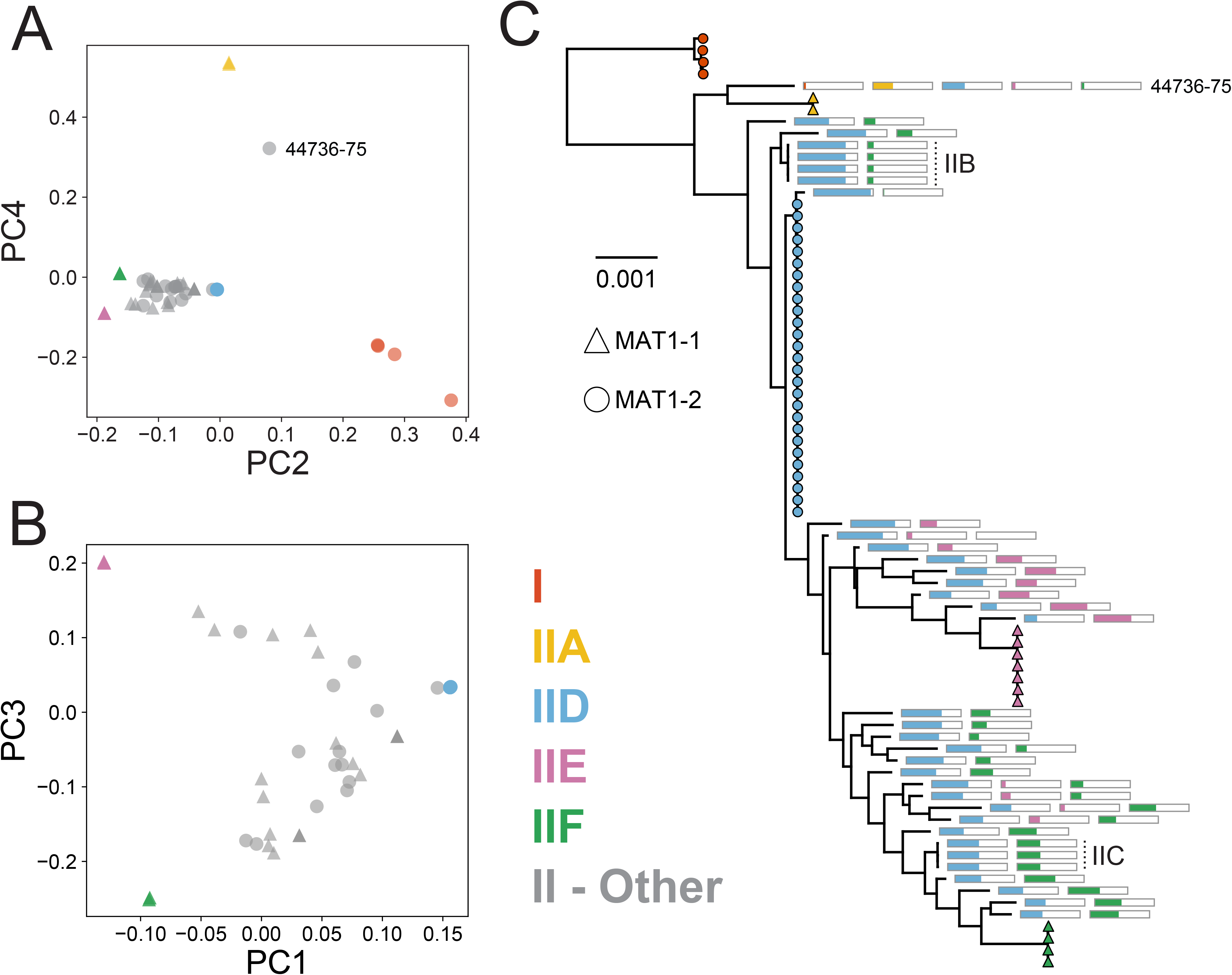
Many North American strains of *Ophidiomyces ophidiicola* are recombinants between clonal lineage IID and clonal lineages IIE and/or IIF. (A) and (B) illustrate the genetic similarity among the strains of Clades I and II based on a co-ancestry matrix generated using ChromoPainter and visualized through principal components (PCs) 1–4. Panel (A) includes all 79 strains, while the strains of Clade I and clonal lineage IIA, as well as strain NWHC 44736-75, have been excluded from panel (B). Shapes indicate the mating type of each strain. Strains from non-recombinant clonal lineages are indicated by colored shapes, recombinant strains are shown in gray. (C) Nuclear maximum-likelihood phylogeny (same as shown in Fig. 1A) with colored bars approximating the level of ancestry from each of the non-recombinant clonal lineages. Strains belonging to non-recombinant clonal lineages are indicated with colored shapes, with the shape indicating mating type. For recombinant strains, each bar represents a set of nuclear single nucleotide polymorphisms (SNPs) that are specific to one of the clonal, non-recombinant lineages, and the proportion of the bar filled with color indicates the proportion of those SNPs that were also observed in that particular strain. A particular bar is only shown if the recombinant strain’s genome contained at least one SNP indicative of that clonal lineage. Colors are the same as those used in (A, B). Vertical dotted lines and labels indicate the recombinant clonal lineages.

We also compared these apparent hybrid strains to their putative parental lineages across all of the variable sites in the nuclear genome, and we found that ≥98% of the SNPs from these putative hybrids can be explained by a combination of the variants observed in lineages IID and IIE or IIF. In fact, in 83% of these strains (29/35), >99.5% of the variable sites can be explained by a combination of two of these clonal lineages. In contrast, <86% of the variable sites can be similarly explained for the strains from lineage IIA, which do not fall along either of these hybridization gradients. Notably, however, several of the hybrid strains appear to share ancestry with lineages IID, IIE, and IIF, which indicates multiple episodes of hybridization (Fig. 2). Repeated hybridization is also consistent with the high variability among recombinants in estimated ancestry from each clonal lineage, as observed in both the PCA analysis (Fig. 2B) and an analysis of fixed SNPs within each clade (Fig. 2C).

We also used PAUP* to look for evidence of homoplasy after excluding the hybrid strains. We observed a substantial increase in the consistency index within Clade II after excluding recent recombinants (i.e., only including lineages IIA and IID–F): 0.93 vs. 0.25 for all of Clade II (Table S3). This is consistent with these recent hybrids being responsible for most of the homoplasy observed within this clade. However, we still observed more homoplasy than we saw within most of the individual clonal lineages, and the consistency index was even lower when we included the four strains from Clade I (0.86). Therefore, despite the absence of recent recombination within each clonal lineage, we did see evidence for historical recombination between lineages.

Geographically, clonal lineage IID was the most widespread in the USA, with strains recovered from 14 eastern states (Fig. 3). Although less frequently sampled, clonal lineage IIE was also widespread, while clonal lineage IIF was more geographically restricted, which could be an artifact of small sample size (IIF was represented by only two strains from wild snakes). Clonal lineage IIA and a related hybrid strain (NWHC 44736-75) were represented by single strains from wild snakes. Geospatial patterns were evident among hybrid strains as well. Specifically, hybrids between lineages IID and IIE were found in the southeastern region of the USA, while hybrids between lineages IID and IIF were found throughout the Atlantic and Gulf Coastal Plains regions. Hybrids of these clonal lineages were rarely sampled in the Midwest region of the USA.

**Fig. 3.**
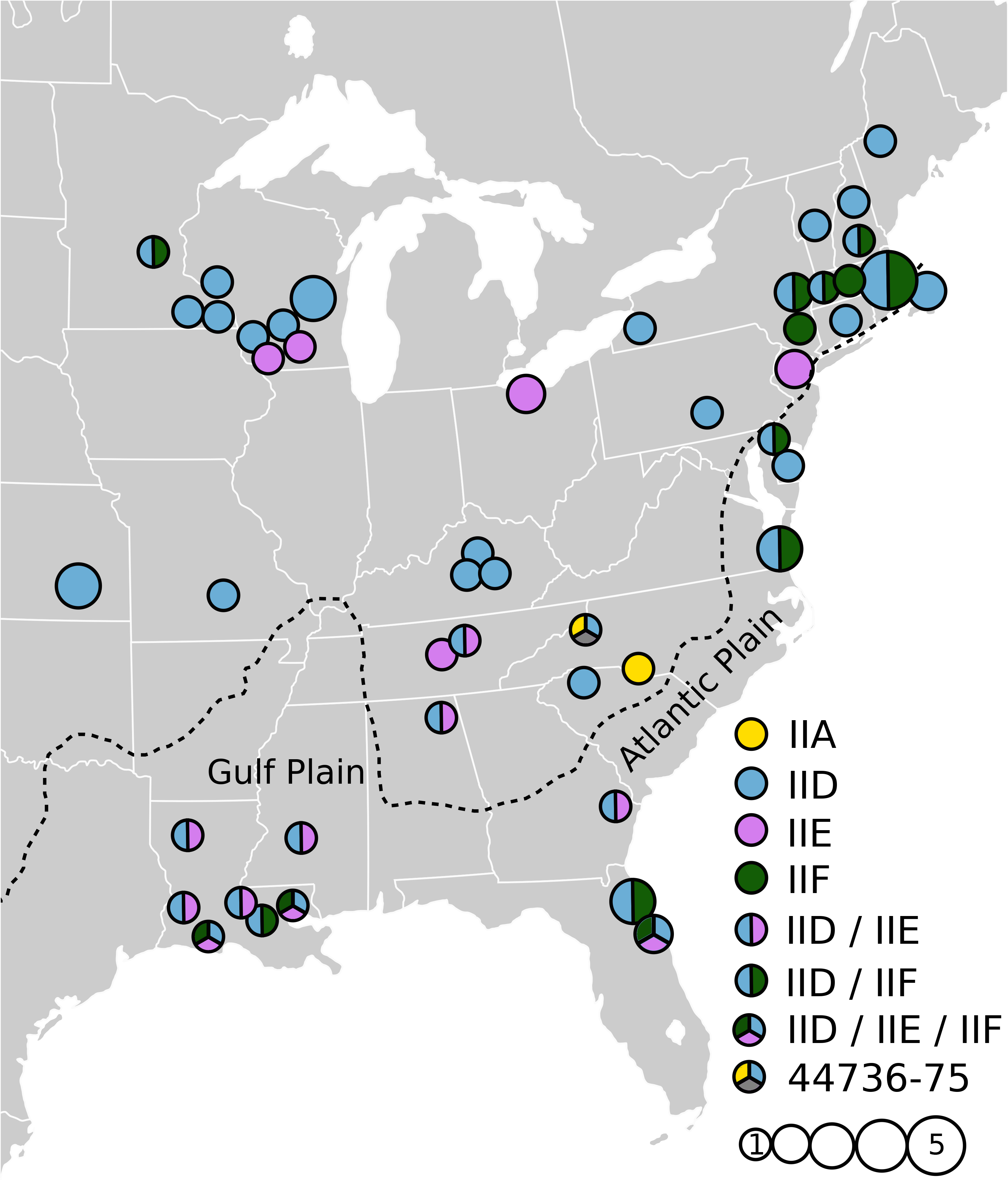
Geographic distribution of *Ophidiomyces ophidiicola* strains isolated from wild snakes in the eastern USA. Solid circles represent non-recombinant clonal lineages. Divided circles depict recombinant strains (color combinations qualitatively indicate genetic signatures of the various lineages present in those recombinants [see Fig. 2]). For strain NWHC 44736-75, a portion of the circle is gray to indicate an unsampled parent lineage. Circle sizes denote the number of strains when multiple strains were collected in proximity to one another. The edge of the Atlantic and Gulf Coastal Plain region is shown with a dashed line.

Within non-recombinant clonal lineages, we examined the relationship between genetic divergence and geographic distance by comparing pairwise nucleotide-level divergence based on the nuclear genome with the distance between localities from which each wild snake was captured. We observed a strong positive correlation between genetic divergence and distance across relatively short geographic distances, but correlation was lacking between these metrics across larger geographic distances (Fig. S3). This pattern is consistent with the presence of fine-scale population structure and an absence of recent long-distance dispersal.

### Molecular Dating Analysis

To assess how long *Oo* has been circulating within the USA, we generated time-structured phylogenies using BEAST. For these analyses, we focused on four subsets of our data, each with a lack of apparent recombination and with evidence for temporal signal as evaluated through comparison of root-to-tip genetic divergence with sampling date (Fig. S4): a mitochondrial analysis that included all 82 strains (Clades I-III; Fig. 4) and nuclear analyses focused independently on the three most commonly sampled clonal lineages: IID–IIF (Fig. 5). Collectively, these three clonal lineages accounted for 52% of the *Oo* strains from wild snakes in the USA and are the parental lineages from which another 45% were derived. Across all analyses, substitution rate estimates were very consistent with the reported rates for the amphibian-infecting fungus *Batrachochytrium dendrobatidis* [16] (Fig. S5). The mean estimated rate for the mitochondrial genome of *Oo* was 9.5 x 10^-7^ (95% highest posterior density [HPD]: 3.6 x 10^-7^ – 1.6 x 10^-6^) substitutions per site per year using the strict clock model. We do not report results for the relaxed clock with the mitochondrial dataset because we did not observe consistent convergence with this model. The mean estimated nuclear substitution rates for *Oo* were 1.9 x 10^-7^ – 1.1 x 10^-6^, including all three clonal lineages and both the strict and relaxed clock models (see Table S4 for details).

**Fig. 4.**
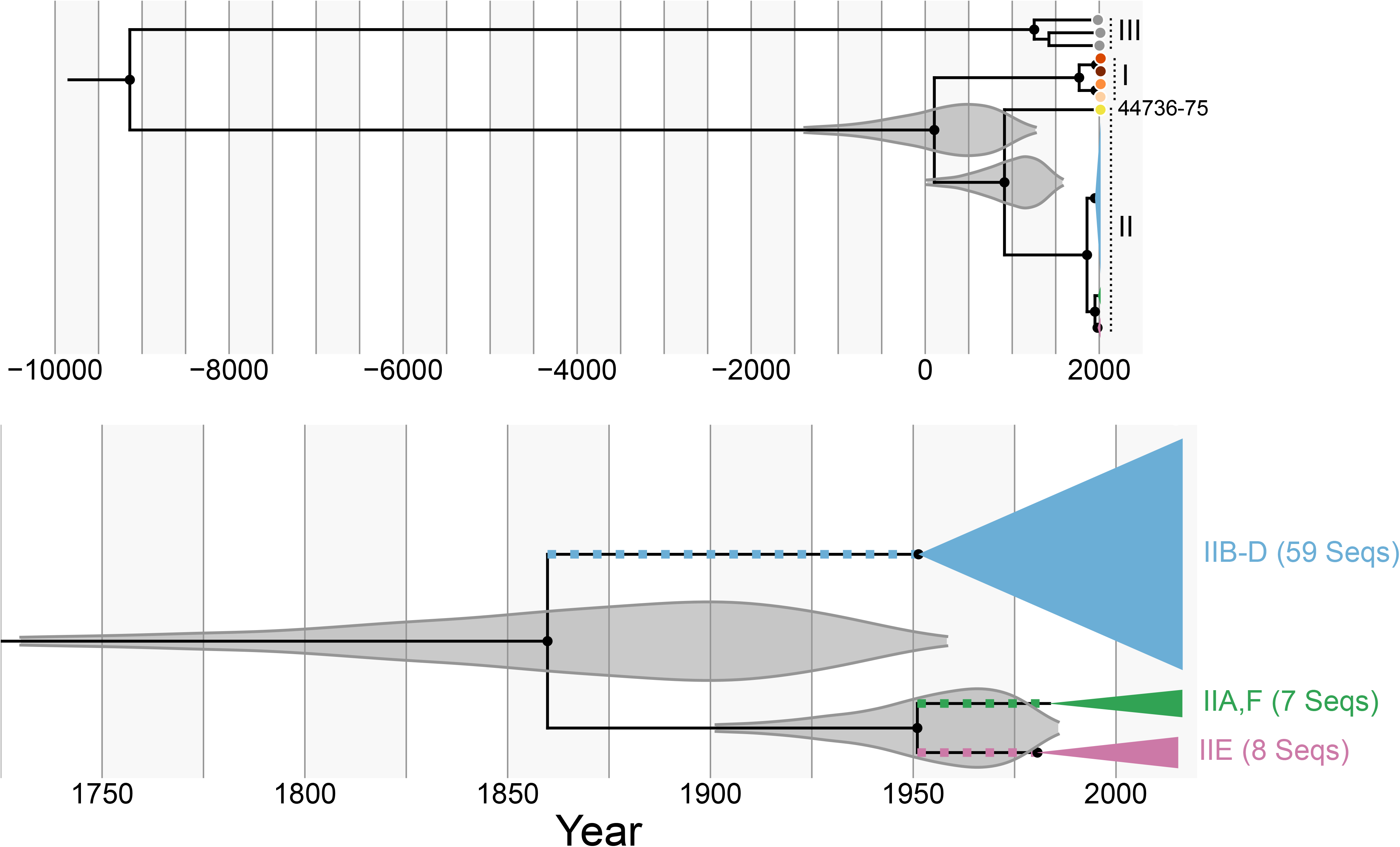
All 79 strains of *Ophidiomyces ophidiicola* from Clades I and II likely shared a common ancestor <4,000 years ago. Time-structured Bayesian phylogeny based on an alignment of 50,624 base pairs of the mitochondrial genome and including all 82 *Oo* strains (see Fig. S1C for an uncollapsed, maximum-likelihood version of the same phylogeny). The top panel shows the entire tree, along with the posterior probability distributions (95% highest posterior density) for the times of the most recent common ancestor (tMRCAs) of Clades I-II and Clade II (gray). The bottom panel shows just a portion of Clade II and the posterior probability distributions (95% highest posterior density) for the tMRCAs of clonal lineages IID-F and IID-E. Colored squares indicate the branches on which we infer that clonal lineages IID-F were first introduced to North America. Nodes represent mean age estimates. Most of the Clade II strains have been collapsed to save space, and the colors and clade/clonal lineage names are identical to those used in Fig. 1. Filled black circles indicate nodes with posterior support ≥95.

**Fig. 5.**
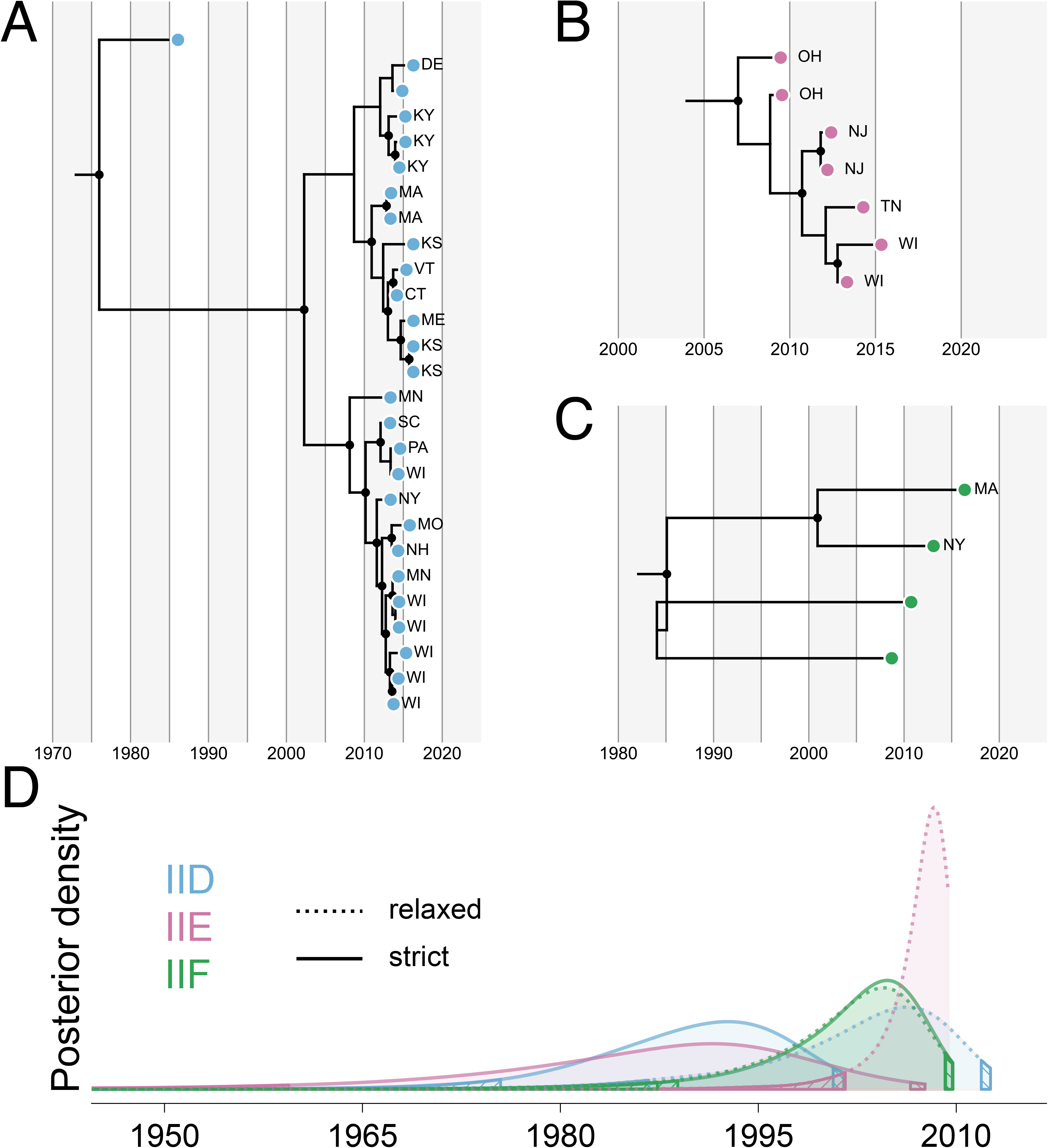
Strains of *Ophidiomyces ophidiicola* within the three most common clonal lineages on wild snakes in North America likely shared a common ancestor within the last ∼50-100 years. (A), (B), and (C) show relaxed clock, time-structured Bayesian phylogenies for Clades IID, IIE, and IIF, respectively. Each phylogeny was built using an alignment of 19,562,499 – 19,658,811 base pairs within the nuclear genome and nodes represent mean age estimates for each node. Filled black circles indicate nodes with posterior support ≥95. For each strain collected from a wild snake, the state within the USA is indicated using two-letter abbreviations (strains from captive snakes are not labeled). (D) Posterior probability distributions for the times of most recent common ancestor (tMRCAs) of clonal lineages IID (blue), IIE (pink), and IIF (green). Estimates from analyses using strict and relaxed clock models are shown with solid and dotted lines, respectively. Only strains isolated from wild snakes were considered in the calculation of tMRCAs. Hatched regions represent the tails of each distribution that fall outside of the 95% highest posterior density.

Based on the mitochondrial analysis, we estimated that Clades I and II diverged ∼2,000 years ago (mean: 110 CE, 95% HPD: 1362 BCE–1267 CE), while Clade III diverged from a common ancestor of Clades I and II ∼11,000 years ago (mean: 9147 BCE, 95% HPD: 17,863–2560 BCE) (Fig. 4). In contrast, we estimated that the vast majority (74/75) of the Clade II strains last shared a common ancestor ∼160 years ago (mean: 1860 CE, 95% HPD: 1731–1958 CE), while the time of the most recent common ancestor (tMRCA) of Clade I likely occurred ∼250 years ago (mean: 1770 CE, 95% HPD: 1553–1938 CE). Within Clade II, strain NWHC 44736-75 was an outlier. The mitochondrial genome from this strain is estimated to have diverged from the rest of Clade II ∼1,100 years ago (tMRCA mean: 914 CE, 95% HPD: 22–1573 CE).

Across both the strict and relaxed clock nuclear analyses, all of the strains from each individual Clade II clonal lineage were estimated to have most likely shared a common ancestor within the last century (Fig. 5, Table S5). For lineages IID and IIF, tMRCA estimates were even more recent when considering only the strains isolated from wild snakes. Mean tMRCA estimates for the isolates from wild snakes from each lineage were 1989/2002, 1985/2007, and 2001/2001 CE for lineages IID, IIE, and IIF, respectively (strict/relaxed clock models), and the oldest date included in any of the 95% HPDs for the wild isolate tMRCAs was 1959 CE (IIE, strict clock analysis). Notably, model testing using both path sampling and stepping stone sampling marginal likelihood estimation approaches found the relaxed clock model to be the best fit for all three clonal lineages (Table S4), and tMRCA estimates for lineages IID and IIE were considerably more recent with the relaxed clock, while the tMRCA estimates for IIF were almost identical with the strict and relaxed models (Table S5).

Our phylogenetic analysis of these individual lineages also revealed several well-supported clades within these lineages, most of which were geographically restricted within our sample set (Fig. 5). Within lineage IIE, for example, our dataset included three pairs of strains, each obtained from wild snakes collected from the same state (New Jersey, Ohio, and Wisconsin). Each pair was phylogenetically closely related and two formed well-supported sublineages within the phylogeny. However, each isolate was distinct from its closest sampled relative, with mean estimated divergence times of ∼1–5 years prior to sampling (based on the relaxed clock analysis).

## DISCUSSION

Phylogenetic and population genetic analyses based on whole genome sequence data can help elucidate the population histories of fungal pathogens and the temporal dynamics of disease emergence. By applying these methods to the causative agent of SFD in the USA, we uncovered a complex history that includes recent expansions of multiple *Oo* clonal lineages as well as recombination between those lineages within a timeframe that is most consistent with the novel pathogen hypothesis [10].

Similar to previous studies that had examined only a small number of loci [26, 27], we identified three major clades among all 82 sampled *Oo* strains, but all strains originating from wild snakes in the USA belonged to a single clade (Clade II). Within this North American clade, we initially observed what appeared to be high genetic diversity as evidenced by the presence of many long phylogenetic branches (Fig. 1A, Fig. S1A). However, after accounting for recombination, we identified just four clonal lineages within which we saw no evidence of recent recombination, although we did observe evidence of potential historical recombination between these clonal lineages (as indicated by some discordance between nuclear and mitochondrial phylogenies, as well as the presence of moderate levels of homoplasy on the branches separating these lineages). Most remaining North American strains not belonging to these four clonal lineages represented recent hybrids between 2-3 of those lineages (Fig. 2). These findings highlight the importance of accounting for recombination when assessing the evolutionary history of fungal pathogen populations.

We identified two mating type loci (*MAT1-1* and *MAT1-2*) among *Oo* strains, consistent with other filamentous ascomycete fungi [36], and both idiomorphs were sampled from wild snakes in the USA at relatively similar frequencies (45% and 55%, respectively). Notably, each clonal lineage within Clade II was characterized by a single mating type and all recombinants involved parent lineages with complementary mating types (i.e., one parent lineage possessing *MAT1-1* locus and one possessing *MAT1-2* locus). Although clonal expansion was observed among recombinant strains in two intensely sampled snake populations (IIB and IIC), most hybrid lineages were sampled only once and had different proportions of their genomes belonging to each parent lineage, indicating unique hybridization events. Furthermore, some strains appeared to be hybrids between more than two clonal lineages (Fig. 2). Altogether, these patterns indicate that hybridization between *Oo* strains has occurred frequently in the USA as a result of sexual reproduction.

In contrast to the endemic pathogen hypothesis, we estimated that all of the North American lineages of *Oo* shared a common ancestor relatively recently. Assuming that there was only a single introduction of *Oo* to the USA, we would predict that this introduction occurred somewhere along the branch separating the MRCA of Clade II with the MRCA of Clades I and II. We estimated that this occurred ∼500–3,400 years ago according to our mitochondrial phylogeny (Fig. 4). The large disparity (four orders of magnitude) between the dates associated with snake movements across Beringia and the genetic divergence of the USA *Oo* lineages makes it unlikely that the fungus was introduced to North America from elsewhere via natural snake movements.

Although the mitochondrial MRCA of the North American clade (Clade II) is estimated to have occurred long before reported cases of SFD (Fig. 4), it is unlikely that this MRCA provides an accurate estimate for the arrival of *Oo* in the USA. The lack of intermediate strains between the main three mitochondrial genotypes observed in the USA (blue, green, and pink in Fig. 1), the absence of non-recombinant intermediates between the nuclear clonal lineages (Fig. 2), and the potentially complex history of recombination involving unsampled lineages for strain NWHC 44736-75 all indicate that these *Oo* lineages did not evolve within North America.

Instead, it is more likely that there have been multiple introductions of *Oo* to North America. In fact, the presence of two mating types indicates that more than one distinct lineage of *Oo* has been introduced to the USA, and the lack of genetic intermediates indicates that each of the non-recombinant clonal lineages (IIA, IID-F), as well as strain NWHC 44736-75, likely represent distinct introductions. Under such a scenario, lineages of *Oo* in the USA would have diverged from a common ancestor that existed outside of North America, and therefore, the MRCA of Clade II would provide an overestimate (i.e., longer ago than is likely realistic) for when the fungus arrived on the continent.

Assuming all three of the primary clonal lineages in the USA (IID-F) represent separate introduction events, our analyses indicate that these introductions likely occurred within the last few hundred years. When excluding strains originating from captive snakes (which were often basal or divergent from strains of the same clonal lineage that were isolated from wild snakes [Fig. 5]), all of the mean tMRCA estimates for these lineages were between 1985 and 2007 (Fig. 5), which is shortly prior to the reported emergence of SFD in wild North American snakes. These tMRCAs likely somewhat underestimate the timing of the initial introductions of these lineages, however, as they only represent the timing of the common ancestors of the sampled strains, and our per lineage sample sizes are relatively small, especially for clonal lineages IIE (n=7 non-recombinant strains from wild snakes) and IIF (n=2). Instead, these should be considered lower bound estimates for the initial introductions of each lineage, while the tMRCAs for IIE-F and IID-F can provide upper bounds for these estimates. Taken together, our molecular clock analyses indicate that these three lineages were likely introduced to North America between 1731 and 2012 for IID and between 1902 and 2009 for IIE and IIF (Fig. 4, Table S5). These ranges are also consistent with a published survey of museum specimens, which demonstrated that at least one lineage of *Oo* was present in the USA as early as 1945 [31].

However, the timing of the tMRCA estimates for the individual clonal lineages (1985–2007) likely does reflect quite recent population expansions within the USA. During an invasion, exotic fungal pathogens often exhibit a lag between the introduction, establishment, and expansion phases [37]. Thus, *Oo* may have resided in the USA for decades before widespread expansion occurred (perhaps facilitated by additional spillover events due to capture and movement of infected wild snakes for the pet trade). Older detections of *Oo* on museum specimens collected from the eastern USA could also represent strains that failed to establish or spread after release.

Tracing *Oo* expansion in the USA is difficult due to a lack of historical isolates and frequent recombination between lineages. However, while some clonal lineages (e.g., lineages IID and IIE) were quite widespread throughout the sampled distribution in the USA, other lineages (i.e., IIA and IIF) and hybrids between certain lineages were more geographically restricted (Fig. 3). Of note, many strains sampled from the Atlantic Coastal Plain were hybrids. This was particularly evident in the Gulf Coastal Plain where all strains were unique recombinants, some involving up to three parent lineages. Such strain diversity may indicate a longer history of contact between the lineages in this region and imply that the initial introduction of at least some *Oo* lineages occurred in the southern USA. Hybridization between lineages is also expected to occur more frequently in regions with high *Oo* prevalence (i.e., more opportunities for co-infections). Thus far, in-depth studies of *Oo* prevalence in snake communities have been very limited in their geographic scope (e.g., [20,38,39]). Future work would be beneficial to understand the relationship between prevalence and the abundance of hybrid strains.

Despite the broad distributions of several clonal lineages, we did not see evidence for recent long-distance dispersal (Fig. S3). Rather, we observed patterns of genetic divergence within clonal lineages that are consistent with fine-scale population structure (Fig. 5). More intensive surveys of *Oo* may reveal spatiotemporal patterns that better elucidate the history of introduction, expansion, and recombination in North America.

In contrast to the recent introductions and subsequent recombination of several clonal lineages in the USA, we identified a single mating type (*MAT1-2*) and no evidence of recombination among our small sample set of *Oo* strains isolated from wild European snakes. The European clade (Clade I) also appears to be older than the North American clonal lineages, with all European strains estimated to have shared an MRCA ∼100-500 years ago. Due to the small sample size (four strains) and geographic coverage (two countries), it is probable that we captured only a small portion of the genetic diversity that exists in Europe; thus, we may have underestimated the time at which the European lineage started expanding. Despite this, the MRCA of the European clade still predates widespread anthropogenic movement of snakes [40].

Introduction of *Oo* to North America could have occurred through transcontinental movement of snakes in captive collections. Indeed, *Oo* strains recovered from captive snakes in Australia, Europe, and North America represented three separate clonal lineages (and hybrids of those lineages) that were also found in wild snakes in the eastern USA, demonstrating that these strains are circulating in captive snake populations across the world. *Oo* has been detected in soil in Malaysia [41], in snakes imported to Russia from Indonesia [42], and in a wild Burmese python (*Python bivittatus*) in Hong Kong [43]. Furthermore, isolates of *Oo* recovered from wild snakes in Taiwan include a representative that belongs to the North American clade (Clade II), as well as a divergent strain most similar to Clade III [27] based on multi-locus sequence typing analysis (Fig. S6). Repeated detections of *Oo* in southeast Asia, along with the presence of such diverse strains of *Oo* in Taiwan, indicate that the fungus may be native to that region [27]. Many fungal pathogens responsible for wildlife epizootics have their origins in Eurasia, including *B. dendrobatidis* [16], *B. salamandrivorans* [44], and *P. destructans* [33]. Most *Oo* detections outside of North America, including those in Eurasia, have occurred through opportunistic sampling; we are unaware of any comprehensive pathogen surveillance or screening efforts targeting *Oo* in other parts of the world. Thus, *Oo* may be much more widely distributed than is currently documented. Therefore, further sampling would be needed to investigate the ‘Out of Eurasia’ hypothesis as the origin of *Oo* in North America.

Although it seems likely that *Oo* was originally introduced to the USA many decades ago, it is possible that the recent emergence of SFD in the eastern USA is linked to subsequent, more recent introductions of novel lineages, geographic expansion of certain clonal lineages, and/or recombination between lineages. Indeed, the range of age estimates for the MRCAs of several of the clonal lineages sampled in the northeastern and midwestern USA is consistent with increased reports of SFD in those regions ([18,21,45]; Fig 5). Due to unknown clinical histories and outcomes for most of the snakes sampled in this study, we were unable to assess whether different *Oo* strains or the presence of particular genes were associated with more severe disease. Controlled studies tracing the fate of wild snakes infected with different strains of *Oo* or challenge experiments in the laboratory would be necessary to determine how pathogenicity varies by lineage, which genes (or combination of genes) might be associated with greater virulence, and whether some hybrid strains possess increased virulence and transmissibility.

Further sampling and strain characterization in other geographic areas will be critical to identify the risk posed by *Oo* to snakes on a global scale. For example, completely naïve snake populations may be most vulnerable to adverse impacts following the introduction of *Oo.* Furthermore, novel contact between previously isolated fungal strains can result in gene transfer and recombination events that may lead to rapid evolution and increased virulence [15, 46], especially when the strains have complementary mating types [47]. For example, if additional sampling supports the presence of a single clonal lineage and mating type (*MAT1-2*) in European snake populations, then those populations could be particularly vulnerable to the introduction of additional strains, especially those possessing the *MAT1-1* locus.

Understanding the mechanisms behind disease emergence often starts with elucidating the history of the causative pathogen. Our data demonstrate recent expansion of *Oo* in North America, likely involving a complex history of multiple strain introductions and recombination that has resulted in a diverse strain pool. While additional work is necessary to determine the exact distribution and origin of *Oo*, differences in strain virulence, and the role of environmental factors in SFD outbreaks, our findings provide critical information for proactive management. Specifically, our data highlight that increased vigilance would be warranted to prevent further spread of *Oo* and the introduction of novel strains to areas where snake populations are at greatest risk.

## MATERIALS AND METHODS

### Ophidiomyces ophidiicola isolates

Eighty-two strains of *Oo* (as determined by internal transcribed spacer region sequence data [48]) were included in this study. All of these strains were obtained from publicly accessible culture collections, isolated at the U.S. Geological Survey - National Wildlife Health Center, or shared by other diagnostic laboratories. These strains represented 65 isolates from wild snakes in the eastern USA, 4 isolates from wild snakes in Europe (n=3 from the United Kingdom; n=1 from Czech Republic), and 13 isolates from captive snakes on three continents (North America, Europe, and Australia). Strains from wild snakes were collected during 2009-2016; those from captive snakes were collected over a broader range of time: 1985-2015. Strains originated from at least 33 snake species and subspecies, representing 18 genera and 6 families of snakes. The list of all strains used in this study, along with associated metadata, are presented in Table S2.

### Whole genome sequencing and annotation

To obtain DNA for whole genome sequencing, cultures of *Oo* were grown in 20 mL of Sabouraud dextrose broth at 24°C for 9-15 days. Fungal biomass from each culture was flash-frozen in liquid nitrogen, lyophilized, pulverized using a pestle and glass beads, resuspended in 700 μ L of LETS buffer (20 mM EDTA [pH 8.0], 0.5% sodium dodecyl sulfate, 10mM Tris-HCl [pH 8.0], 0.1 M LiCl), and DNA was extracted using the phenol-chloroform method.

Library preparation and whole genome sequencing were performed by the University of Wisconsin-Madison Biotechnology Center (Madison, Wisconsin). Seven strains (NWHC 22687-1, NWHC 23942-1, NWHC 24266-2, NWHC 45692-2, CBS 122913 [type isolate of *Oo*], UAMH 10296, UAMH 9985) were sequenced on an Illumina MiSeq system (2 x 250 bp read lengths). The remaining 75 strains were sequenced on an Illumina HiSeq 2500 sequencing system using a high-output kit and 2 x 125 bp read lengths.

Nuclear and mitochondrial *de novo* assemblies were generated using the Automatic Assembly for the Fungi (AAFTF) pipeline [49]. Briefly, the AAFTF pipeline ran the following steps: the Illumina data were cleaned (i.e. removal of adapters, artifacts, contaminants, etc.) using BBDuk from the BBMAP v38.92 package [50], the mitochondrial genome was assembled using NOVOplasty v4.2 [51], the nuclear genome was assembled with SPAdes v3.10.1 [52], and the resulting nuclear assembly was error corrected with three polishing rounds using Pilon v1.24 [52, 53]. Assembled genomes were deposited in the National Center for Biotechnology Information (NCBI) database under BioProject PRJNA780910 (see Table S2).

Genome annotation was completed using funannotate v1.5.2 [54]. A library of repetitive sequences found in *Oo* was generated for the type strain (CBS 122913) using the funannotate wrapper for RepeatModeler v1.0.11 [55] and RepeatMasker v4.0.7 [56]. The resulting library was used to soft-mask all of the *de novo* assemblies in this study using the ‘funannotate mask’ command. Subsequently *de novo* gene prediction training parameters for AUGUSTUS v3.2.1 [57] were generated from strain CBS 122913 by utilizing the automated training pipeline in ‘funannotate predict.’ Briefly, BUSCO [58] was used to generate initial gene predictions used for training AUGUSTUS. The AUGUSTUS training parameters were used on all genomes in this study. The ‘funannotate predict’ pipeline also utilized gene predictions from GeneMark v4.32 [59] to compute consensus gene model predictions using EvidenceModeler [60].

### Mating type

To identify the mating type idiomorph in whole genome sequence datasets, we aligned the Illumina reads for each strain to the CBS 122913 genome assembly using minimap2 v2.12 [61]. We then calculated the ratio of read mapping coverage at the *MAT1-1* locus (scaffold_5:201703-202676) compared to a region outside of the mating type idiomorph (scaffold_5:196703-197676) using BEDTools [62]. To identify *MAT1-2*, we aligned the conserved flanking regions of *MAT1-1* from strain CBS 122913 with those of a strain in which *MAT1-1* was not detected and identified predicted genes between those flanking regions. To further investigate which *MAT1* loci were present in a larger sample set from Europe, we screened nine additional strains of *Oo* isolated from wild snakes in the United Kingdom for which whole-genome sequencing was not performed for the presence of *MAT1-1* and *MAT1-2* using two newly designed PCR assays (see Supporting Text; Table S2).

### Phylogenetic Analysis

We used Proteinortho v6 [63] to identify putatively orthologous nuclear protein-coding genes across all 82 *Oo* strains using amino acid sequences as input. We individually aligned amino acid sequences for each of the 5,811 single copy genes present in all 82 isolates using MAFFT v7.487 with default settings [64], and then concatenated all of these aligned proteins, which resulted in a single amino acid-level, genome-wide alignment that consisted of 3,311,400 positions. Using this alignment, we generated a maximum-likelihood phylogeny using RAxML v8.2.12 [65]. We used RAxML’s automatic protein model assignment algorithm (-m PROTGAMMAAUTO), which selected the JTT model as the best fit, along with 100 rapid bootstrapping iterations followed by a maximum-likelihood search (-f a -N 100).

To examine mitochondrial relationships among strains, we identified SNPs using NASP v1.2.0 [66] with BWA-mem v0.7.7 and GATK v3.4-46-gbc02625. As a reference, we used our *de novo* assembly for the type strain of *Oo*, CBS 122913, and we utilized all reference positions at which a high-confidence call was made for ≥ 80% of the strains. Using the same approach, we also identified nuclear SNPs for the 79 isolates of *Oo* from Clades I and II. For this analysis, we utilized 38 scaffolds previously assembled from the MYCO-ARIZ AN0400001 strain of *Oo* as a reference (GenBank: MWKM01000001.1 - MWKM01000021.1, MWKM01000023.1 - MWKM01000039.1). Prior to running NASP, we removed sequencing adapters and trimmed low quality bases using BBDuk (last modified 2019-01-23) [50]. We then generated maximum-likelihood phylogenies using RAxML-NG v0.5.1b [67] with the GTR+G model, 10 randomized parsimony starting trees, and 100 bootstrap replicates. For the mitochondrial SNP alignment, we also constructed a median-joining haplotype network [68] using PopART [69].

### Recombination Analyses

We used PAUP* v4.0a (build 168) to build parsimony-based phylogenies and to calculate consistency/homoplasy indexes. Trees were constructed using random stepwise addition with 100 replicates and tree-bisection-reconnection with a reconnection limit of 8. PhiPack [70] was used to examine recombination using the Phi test [34]. Analyses were performed individually for each reference scaffold (using all positions with high confidence calls in ≥80% of the 75 strains from Clade II) with a window size of 50,000 bases and a step size of 25,000 bases.

To generate a co-ancestry matrix for the 79 strains from Clades I and II, we used ChromoPainter within FineSTRUCTURE v4.1.1 [35]. We used the default parameters implemented via the ‘automatic’ mode, with ploidy set to 1 and a fixed recombination rate of 0.000001 Morgans. We then conducted a PCA using the “chunkcounts” file generated by ChromoPainter along with the *mypca* function contained in the FineSTRUCTURE R library [71].

The capture locations for most snakes were available only to the county level. Thus, we used county centroids (or state centroids when state was the only locality information available) as coordinates to represent origin locations for *Oo* strains. Distances between strain collection localities were calculated with the Haversine formula using the geosphere package [72] in R [73].

### Molecular Dating Analysis

To estimate dates of divergence for the major *Oo* clades, we used BEAST v1.10.5 with BEAGLE v.3.2.0 [74]. For the mitochondrial genome, we conducted a single analysis including all 82 strains and all reference positions at which a genotype was called in ≥80% of the strains (50,624 bp, 817 variable sites). Although we observed some discordance between nuclear and mitochondrial phylogenies within Clade II, we did not observe any discordance in clade membership, and therefore analysis of the mitochondrial genome can be used to date the divergence between clades. Because of a history of recombination in the nuclear genome, we ran separate analyses for each of the three primary clonal lineages in Clade II (IID-IIF) using all reference positions at which a genotype was called for 100% of the included strains (19,562,499– 19,658,811 bp, 324–1,635 variable sites). For the nuclear datasets, we used ClonalFrameML [75] to identify and remove any regions with evidence of recombination. This analysis only flagged 16, 41, and 0 bp as potentially recombinant for clonal lineages IID, IIE and IIF, respectively.

All four analyses used the general time reversible (GTR) substitution model including invariant sites and a discrete gamma distribution of rates among sites with four categories and a coalescent tree model assuming constant population size over time [76] with a lognormal prior distribution (*μ*=0, *σ*=1, offset=0). For each dataset, we compared two clock models: a strict clock and an uncorrelated relaxed clock model with a lognormal distribution, both with a continuous-time Markov chain rate reference prior [77]. Each analysis was run with a chain length of ≥500M and parameters were logged 10,000 times per run, at regular intervals. Each dataset and model combination was run at least twice with different random seeds to ensure consistent convergence. We also tested additional tree models (e.g., Bayesian Skyline and Bayesian SkyGrid); however, none of our datasets reached convergence with these more complex tree priors.

## DATA AVAILABILITY

Metadata associated with this project are available at https://doi.org/10.5066/P9J2MCLJ. Additional code and output for various analyses have been deposited in Open Science Forum and are available at https://osf.io/evzk3/. Sequence data, genome assemblies, and genome annotations associated with this project were deposited in GenBank under BioProject PRJNA780910.

## ACKNOWLEDGEMENTS

We thank the numerous state and federal biologists, contractors, wildlife health staff, scientists, and volunteers, including personnel at the U.S. Geological Survey - National Wildlife Health Center, that participated in collecting and compiling metadata, as well as collecting and processing samples that led to the isolation of *Ophidiomyces ophidiicola* strains from the USA. We also thank the Garden Wildlife Health project (www.gardenwildlifehealth.org, a wildlife disease surveillance program coordinated by the Institute of Zoology, UK), Angela Winnett, Dr. Iain Barr, University of East Anglia, UK, and Dr. Dave Leech, British Trust for Ornithology, UK, and Dr. Vojtech Baláž, University of Veterinary and Sciences Brno who similarly assisted with collection of metadata and samples in Europe. We are grateful to the various laboratories that preserved and shared strains that were previously published on, including the Cornell University College of Veterinary Medicine (Dr. Krysten Schuler), the Southeastern Cooperative Wildlife Disease Study, and the University of Florida College of Veterinary Medicine (Dr. Jim Wellehan). This project was funded by the U.S. Geological Survey. J.T.L. was supported by the State of Arizona Technology and Research Initiative Fund (TRIF, administered by the Arizona Board of Regents, through Northern Arizona University). Any use of trade, firm, or product names is for descriptive purposes only and does not imply endorsement by the U.S. Government.

The authors declare no competing interests.

## REFERENCES

1. Harvell CD, Kim K, Burkholder JM, Colwell RR, Epstein PR, Grimes DJ, et al. Emerging marine diseases--climate links and anthropogenic factors. Science. 1999;285: 1505–1510.

2. Daszak P, Cunningham AA, Hyatt AD. Emerging infectious diseases of wildlife--threats to biodiversity and human health. Science. 2000;287: 443–449.

3. Dobson A, Foufopoulos J. Emerging infectious pathogens of wildlife. Philos Trans R Soc Lond B Biol Sci. 2001;356: 1001–1012.

4. Tompkins DM, Carver S, Jones ME, Krkošek M, Skerratt LF. Emerging infectious diseases of wildlife: a critical perspective. Trends Parasitol. 2015;31: 149–159.

5. Fisher MC, Gow NAR, Gurr SJ. Tackling emerging fungal threats to animal health, food security and ecosystem resilience. Philos Trans R Soc Lond B Biol Sci. 2016;371. doi:10.1098/rstb.2016.0332

6. Cunningham AA, Daszak P, Wood JLN. One Health, emerging infectious diseases and wildlife: two decades of progress? Philos Trans R Soc Lond B Biol Sci. 2017;372. doi:10.1098/rstb.2016.0167

7. Fisher MC, Henk DA, Briggs CJ, Brownstein JS, Madoff LC, McCraw SL, et al. Emerging fungal threats to animal, plant and ecosystem health. Nature. 2012;484: 186–194.

8. Cheng TL, Reichard JD, Coleman JTH, Weller TJ, Thogmartin WE, Reichert BE, et al. The scope and severity of white-nose syndrome on hibernating bats in North America. Conserv Biol. 2021;35: 1586–1597.

9. Scheele BC, Pasmans F, Skerratt LF, Berger L, Martel A, Beukema W, et al. Amphibian fungal panzootic causes catastrophic and ongoing loss of biodiversity. Science. 2019;363: 1459–1463.

10. Rachowicz LJ, Hero J-M, Alford RA, Taylor JW, Morgan JAT, Vredenburg VT, et al. The Novel and Endemic Pathogen Hypotheses: Competing Explanations for the Origin of Emerging Infectious Diseases of Wildlife. Conservation Biology. 2005. pp. 1441–1448. doi:10.1111/j.1523-1739.2005.00255.x

11. Juzwik J, Harrington TC, MacDonald WL, Appel DN. The origin of Ceratocystis fagacearum, the oak wilt fungus. Annu Rev Phytopathol. 2008;46: 13–26.

12. Broders KD, Boraks A, Sanchez AM, Boland GJ. Population structure of the butternut canker fungus, Ophiognomonia clavigignenti-juglandacearum, in North American forests. Ecol Evol. 2012;2: 2114–2127.

13. Grünwald NJ, Garbelotto M, Goss EM, Heungens K, Prospero S. Emergence of the sudden oak death pathogen *Phytophthora ramorum*. Trends Microbiol. 2012;20: 131–138.

14. Desprez-Loustau M-L, Robin C, Buée M, Courtecuisse R, Garbaye J, Suffert F, et al. The fungal dimension of biological invasions. Trends Ecol Evol. 2007;22: 472–480.

15. Farrer RA, Weinert LA, Bielby J, Garner TWJ, Balloux F, Clare F, et al. Multiple emergences of genetically diverse amphibian-infecting chytrids include a globalized hypervirulent recombinant lineage. Proc Natl Acad Sci U S A. 2011;108: 18732–18736.

16. O’Hanlon SJ, Rieux A, Farrer RA, Rosa GM, Waldman B, Bataille A, et al. Recent Asian origin of chytrid fungi causing global amphibian declines. Science. 2018;360: 621–627.

17. Sutherland WJ, Aveling R, Brooks TM, Clout M, Dicks LV, Fellman L, et al. A horizon scan of global conservation issues for 2014. Trends Ecol Evol. 2014;29: 15–22.

18. Lorch JM, Knowles S, Lankton JS, Michell K, Edwards JL, Kapfer JM, et al. Snake fungal disease: an emerging threat to wild snakes. Philos Trans R Soc Lond B Biol Sci. 2016;371. doi:10.1098/rstb.2015.0457

19. Funk S, Bogich TL, Jones KE, Kilpatrick AM, Daszak P. Quantifying trends in disease impact to produce a consistent and reproducible definition of an emerging infectious disease. PLoS One. 2013;8: e69951.

20. Davy CM, Shirose L, Campbell D, Dillon R, McKenzie C, Nemeth N, et al. Revisiting ophidiomycosis (snake fungal disease) after a decade of targeted research. Front Vet Sci. 2021;8: 665805.

21. Allender MC, Dreslik M, Wylie S, Phillips C, Wylie DB, Maddox C, et al. *Chrysosporium* sp. infection in eastern massasauga rattlesnakes. Emerg Infect Dis. 2011;17: 2383–2384.

22. Allender MC, Raudabaugh DB, Gleason FH, Miller AN. The natural history, ecology, and epidemiology of *Ophidiomyces ophiodiicola* and its potential impact on free-ranging snake populations. Fungal Ecology. 2015. pp. 187–196. doi:10.1016/j.funeco.2015.05.003

23. Lorch JM, Lankton J, Werner K, Falendysz EA, McCurley K, Blehert DS. Experimental infection of snakes with *Ophidiomyces ophiodiicola* causes pathological changes that typify snake fungal disease. MBio. 2015;6: e01534–15.

24. Allender MC, Baker S, Wylie D, Loper D, Dreslik MJ, Phillips CA, et al. Development of snake fungal disease after experimental challenge with *Ophidiomyces ophiodiicola* in cottonmouths (*Agkistrodon piscivorous*). PLoS One. 2015;10: e0140193.

25. Burbrink FT, Lorch JM, Lips KR. Host susceptibility to snake fungal disease is highly dispersed across phylogenetic and functional trait space. Sci Adv. 2017;3: e1701387.

26. Franklinos LHV, Lorch JM, Bohuski E, Rodriguez-Ramos Fernandez J, Wright ON, Fitzpatrick L, et al. Emerging fungal pathogen *Ophidiomyces ophiodiicola* in wild European snakes. Sci Rep. 2017;7: 3844.

27. Sun P-L, Yang C-K, Li W-T, Lai W-Y, Fan Y-C, Huang H-C, et al. Infection with *Nannizziopsis guarroi* and *Ophidiomyces ophiodiicola* in reptiles in Taiwan. Transbound Emerg Dis. 2021. doi:10.1111/tbed.14049

28. Campbell LJ, Burger J, Zappalorti RT, Bunnell JF, Winzeler ME, Taylor DR, et al. Soil reservoir dynamics of *Ophidiomyces ophidiicola*, the causative agent of snake fungal disease. Journal of Fungi. 2021. p. 461. doi:10.3390/jof7060461

29. Townsend TM, Leavitt DH, Reeder TW. Intercontinental dispersal by a microendemic burrowing reptile (Dibamidae). Proc Biol Sci. 2011;278: 2568–2574.

30. Guo P, Liu Q, Xu Y, Jiang K, Hou M, Ding L, et al. Out of Asia: Natricine snakes support the Cenozoic Beringian Dispersal Hypothesis. Mol Phylogenet Evol. 2012;63: 825–833.

31. Lorch JM, Price SJ, Lankton JS, Drayer AN. Confirmed cases of ophidiomycosis in museum specimens from as early as 1945, United States. Emerg Infect Dis. 2021;27: 1986– 1989.

32. Grünwald NJ, Goss EM. Evolution and population genetics of exotic and re-emerging pathogens: Novel tools and approaches. Annu Rev Phytopathol. 2011;49: 249–267.

33. Drees KP, Lorch JM, Puechmaille SJ, Parise KL, Wibbelt G, Hoyt JR, et al. Phylogenetics of a fungal invasion: Origins and widespread dispersal of white-nose syndrome. MBio. 2017;8. doi:10.1128/mBio.01941-17

34. Bruen TC, Philippe H, Bryant D. A simple and robust statistical test for detecting the presence of recombination. Genetics. 2006. pp. 2665–2681. doi:10.1534/genetics.105.048975

35. Lawson DJ, Hellenthal G, Myers S, Falush D. Inference of population structure using dense haplotype data. PLoS Genet. 2012;8: e1002453.

36. Turgeon BG, Yoder OC. Proposed nomenclature for mating type genes of filamentous ascomycetes. Fungal Genet Biol. 2000;31: 1–5.

37. Garbelotto M. Molecular analysis to study invasions by forest pathogens: examples from the Mediterranean ecosystems. Phytopathol Mediterr. 2008;47: 183–203.

38. McKenzie JM, Price SJ, Fleckenstein JL, Drayer AN, Connette GM, Bohuski E, et al. Field diagnostics and seasonality of *Ophidiomyces ophiodiicola* in wild snake populations. Ecohealth. 2019;16: 141–150.

39. Haynes E, Chandler HC, Stegenga BS, Adamovicz L, Ospina E, Zerpa-Catanho D, et al. Ophidiomycosis surveillance of snakes in Georgia, USA reveals new host species and taxonomic associations with disease. Sci Rep. 2020;10: 10870.

40. Hierink F, Bolon I, Durso AM, de Castañeda RR, Zambrana-Torrelio C, Eskew EA, et al. Forty-four years of global trade in CITES-listed snakes: Trends and implications for conservation and public health. Biological Conservation. 2020. p. 108601. doi:10.1016/j.biocon.2020.108601

41. Ali SRA, Safari S, Thakib MS, Bakeri SA, Ghani NAA. Soil fungal community associated with peat in Sarawak identified using 18S rDNA marker. Journal of Palm Oil Research. 2016. pp. 161–171. doi:10.21894/jopr.2016.2802.03

42. Ovchinnikov RS, Vasilyev DB, Gaynullina AG, Yuzhakov AG, Kapustin AV, Savinov VA, et al. Detection of *Ophidiomyces ophidiicola* in three file snakes (Acrochordus granulatus) imported from Indonesia to the Moscow Zoo (Russia). Journal of Zoo and Wildlife Medicine. 2021. doi:10.1638/2020-0091

43. Grioni A, To KW, Crow P, Rose-Jeffreys L, Ching KK, Chi LO, et al. Detection of *Ophidiomyces ophidiicola* in a wild Burmese python (*Python bivittatus*) in Hong Kong SAR, China. J Herpetol Med Surg. 2021;31: 283–291.

44. Martel A, Blooi M, Adriaensen C, Van Rooij P, Beukema W, Fisher MC, et al. Wildlife disease. Recent introduction of a chytrid fungus endangers Western Palearctic salamanders. Science. 2014;346: 630–631.

45. Allender MC, Phillips CA, Baker SJ, Wylie DB, Narotsky A, Dreslik MJ. Hematology in an eastern massasauga (*Sistrurus catenatus*) population and the emergence of *Ophidiomyces* in Illinois, USA. J Wildl Dis. 2016;52: 258–269.

46. Stukenbrock EH, McDonald BA. The origins of plant pathogens in agro-ecosystems. Annu Rev Phytopathol. 2008;46: 75–100.

47. Brasier CM, Kirk SA. Rapid emergence of hybrids between the two subspecies of *Ophiostoma novo-ulmi* with a high level of pathogenic fitness. Plant Pathol. 2010;59: 186– 199.

48. Sigler L, Hambleton S, Paré JA. Molecular characterization of reptile pathogens currently known as members of the *Chrysosporium* anamorph of *Nannizziopsis vriesii* complex and relationship with some human-associated isolates. J Clin Microbiol. 2013;51: 3338–3357.

49. Stajich J, Palmer J. AAFTF: Automatic Assembly for the Fungi. 2018. Available: https://github.com/stajichlab/AAFTF

50. Bushnell B. BBMap. 2014. Available: https://sourceforge.net/projects/bbmap/files/

51. Dierckxsens N, Mardulyn P, Smits G. NOVOPlasty: *de novo* assembly of organelle genomes from whole genome data. Nucleic Acids Research. 2016. p. gkw955. doi:10.1093/nar/gkw955

52. Bankevich A, Nurk S, Antipov D, Gurevich AA, Dvorkin M, Kulikov AS, et al. SPAdes: A new genome assembly algorithm and its applications to single-cell sequencing. J Comput Biol. 2012;19: 455–477.

53. Walker BJ, Abeel T, Shea T, Priest M, Abouelliel A, Sakthikumar S, et al. Pilon: An integrated tool for comprehensive microbial variant detection and genome assembly improvement. PLoS One. 2014;9: e112963.

54. Palmer J, Stajich JE. Funannotate: A fungal genome annotation and comparative genomics pipeline. 2016. doi:10.5281/zenodo3354704

55. Smit AFA, Hubley R. RepeatModeler. 2017. Available: http://www.repeatmasker.org

56. Smit AFA, Hubley R, Green P. RepeatMasker. 2017. Available: http://www.repeatmasker.org

57. Stanke M, Morgenstern B. AUGUSTUS: A web server for gene prediction in eukaryotes that allows user-defined constraints. Nucleic Acids Research. 2005. pp. W465–W467. doi:10.1093/nar/gki458

58. Simão FA, Waterhouse RM, Ioannidis P, Kriventseva EV, Zdobnov EM. BUSCO: Assessing genome assembly and annotation completeness with single-copy orthologs. Bioinformatics. 2015;31: 3210–3212.

59. Besemer J, Borodovsky M. GeneMark: Web software for gene finding in prokaryotes, eukaryotes and viruses. Nucleic Acids Res. 2005;33: W451–4.

60. Haas BJ, Salzberg SL, Zhu W, Pertea M, Allen JE, Orvis J, et al. Automated eukaryotic gene structure annotation using EVidenceModeler and the Program to Assemble Spliced Alignments. Genome Biol. 2008;9: R7.

61. Li H. Minimap2: Pairwise alignment for nucleotide sequences. Bioinformatics. 2018;34: 3094–3100.

62. Quinlan AR, Hall IM. BEDTools: A flexible suite of utilities for comparing genomic features. Bioinformatics. 2010;26: 841–842.

63. Lechner M, Findeiss S, Steiner L, Marz M, Stadler PF, Prohaska SJ. Proteinortho: Detection of (co-)orthologs in large-scale analysis. BMC Bioinformatics. 2011;12: 124.

64. Katoh K, Standley DM. MAFFT multiple sequence alignment software version 7: Improvements in performance and usability. Mol Biol Evol. 2013;30: 772–780.

65. Stamatakis A. RAxML version 8: A tool for phylogenetic analysis and post-analysis of large phylogenies. Bioinformatics. 2014;30: 1312–1313.

66. Sahl JW, Lemmer D, Travis J, Schupp JM, Gillece JD, Aziz M, et al. NASP: An accurate, rapid method for the identification of SNPs in WGS datasets that supports flexible input and output formats. Microb Genom. 2016;2: e000074.

67. Kozlov AM, Darriba D, Flouri T, Morel B, Stamatakis A. RAxML-NG: A fast, scalable and user-friendly tool for maximum likelihood phylogenetic inference. Bioinformatics. 2019;35: 4453–4455.

68. Bandelt HJ, Forster P, Röhl A. Median-joining networks for inferring intraspecific phylogenies. Mol Biol Evol. 1999;16: 37–48.

69. Leigh JW, Bryant D. popart: Full-feature software for haplotype network construction. Methods in Ecology and Evolution. 2015. pp. 1110–1116. doi:10.1111/2041-210x.12410

70. Bruen T. PhiPack: PHI test and other tests of recombination. [cited 15 Oct 2021]. Available: https://www.maths.otago.ac.nz/~dbryant/software/phimanual.pdf

71. Lawson D. In: PaintMyChromosomes.com fineSTRUCTURE v2 & GLOBETROTTER [Internet]. 21 May 2018 [cited 15 Oct 2021]. Available: http://www.paintmychromosomes.com/

72. Hijmans RJ. Geosphere: Spherical trigonometry. 2021. Available: https://CRAN.R-project.org/package=geosphere

73. R Core Team. R: A language and environment for statistical computing. 2021.

74. Suchard MA, Lemey P, Baele G, Ayres DL, Drummond AJ, Rambaut A. Bayesian phylogenetic and phylodynamic data integration using BEAST 1.10. Virus Evol. 2018;4: vey016.

75. Didelot X, Wilson DJ. ClonalFrameML: Efficient inference of recombination in whole bacterial genomes. PLoS Comput Biol. 2015;11: e1004041.

76. Drummond AJ, Nicholls GK, Rodrigo AG, Solomon W. Estimating mutation parameters, population history and genealogy simultaneously from temporally spaced sequence data. Genetics. 2002;161: 1307–1320.

77. Ferreira MAR, Suchard MA. Bayesian analysis of elapsed times in continuous-time Markov chains. Canadian Journal of Statistics. 2008. pp. 355–368. doi:10.1002/cjs.5550360302

