## Supporting Text for "Population genetic analysis of *Ophidiomyces ophidiicola*, the causative agent of snake fungal disease, indicates recent introductions to the USA"

This file includes:

Fig. S1

Fig. S2

Fig. S3

Fig. S4

Fig. S5

Fig. S6

Table S1

Table S2 legend

Table S3

Table S4 legend

Table S5

Supporting Text

Supporting References

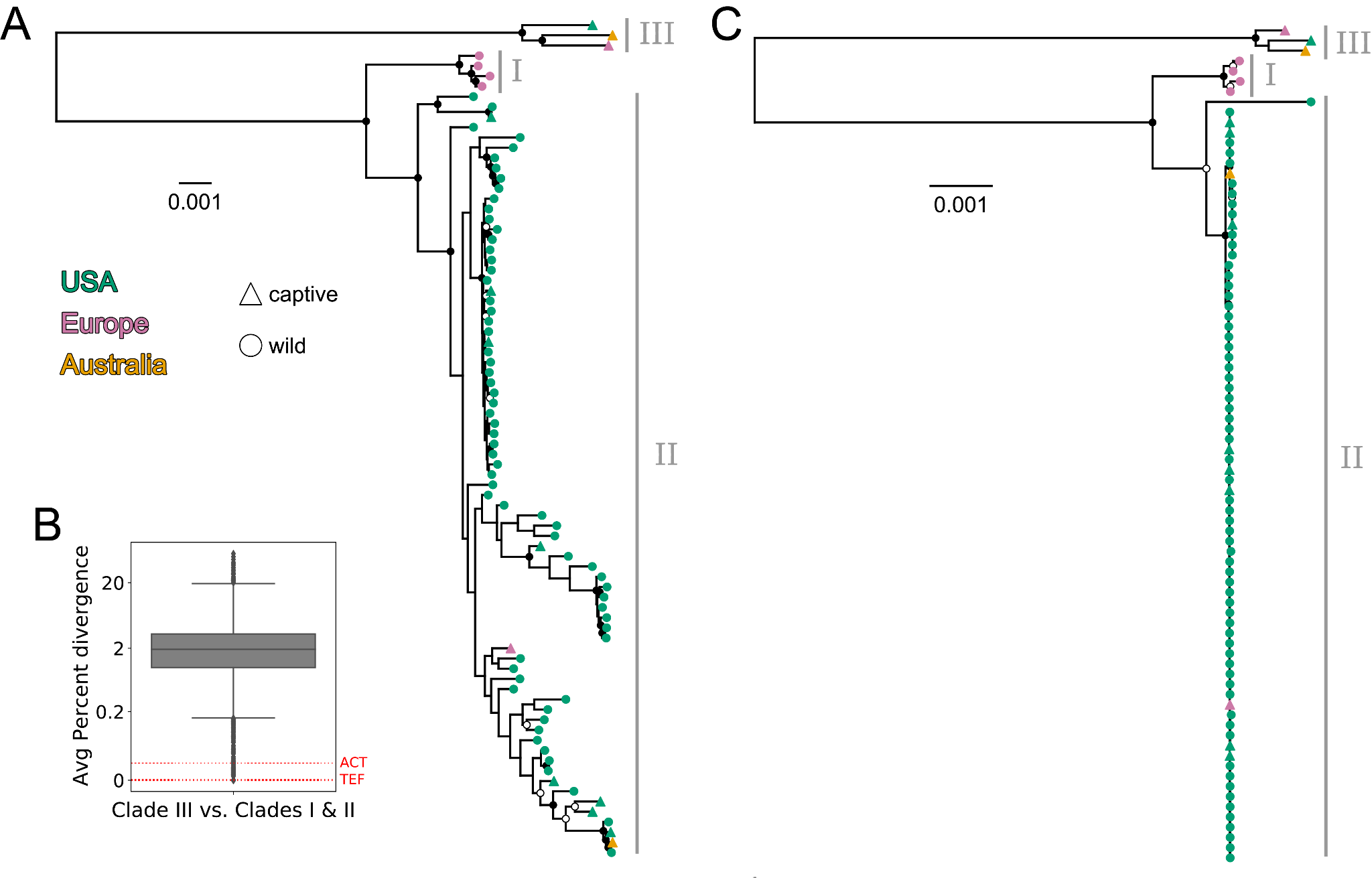

**Fig. S1. All North American *Ophidiomyces ophidiicola* (*Oo*) strains isolated from wild snakes belong to a distinct phylogenetic clade.** (A) Maximum-likelihood phylogeny including all 82 *Oo* strains and based on an amino acid-level alignment consisting of 5,811 nuclear proteins and 3,311,400 positions. (B) Boxplot showing the mean, pairwise percent divergence between Clade III and Clades I and II across the 5,811 proteins included in the alignment for (A). The red dashed lines indicate the values for the two protein coding genes that have been used previously in phylogenetic studies of *Oo*: actin (*ACT*) and translation elongation factor 2ɑ (*TEF*). The limits of the box correspond to the 1st and 3rd quartiles, the black line inside the box corresponds to the median, and the whiskers extend to points that lie within 1.5 interquartile ranges of the 1st and 3rd quartiles. (C) Maximum-likelihood phylogeny including all 82 *Oo* strains and based on a reference-based, nucleotide-level mitochondrial alignment (50,624 positions). In both (A) and (C), tip shapes indicate whether the infected snake was in captivity (triangles) or wild (circles). Filled black and white circles indicate nodes with bootstrap support ≥90 and ≥70, respectively. Gray vertical lines and labels indicate the three primary clades.

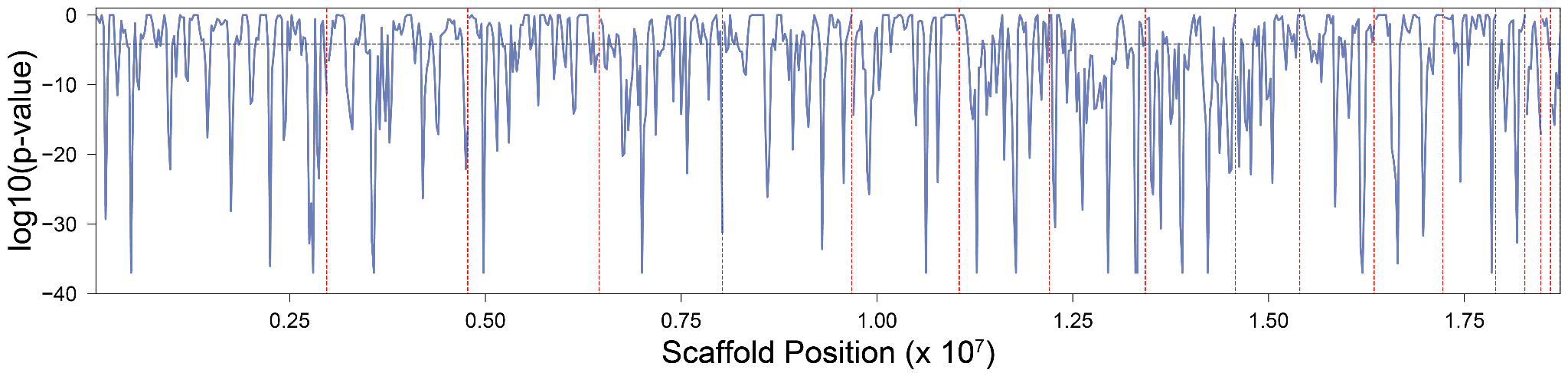
**Fig. S2. Evidence for recombination within Clade II throughout the *Ophidiomyces ophidiicola* genome.** Log-transformed Phi test p-values calculated using PhiPack with a window size of 50,000 bases and a step size of 25,000 bases (1). Black horizontal dashed line indicates a p-value of 0.05 with Bonferroni correction for 749 different tests. Red vertical dashed lines indicate scaffold boundaries within the reference assembly.

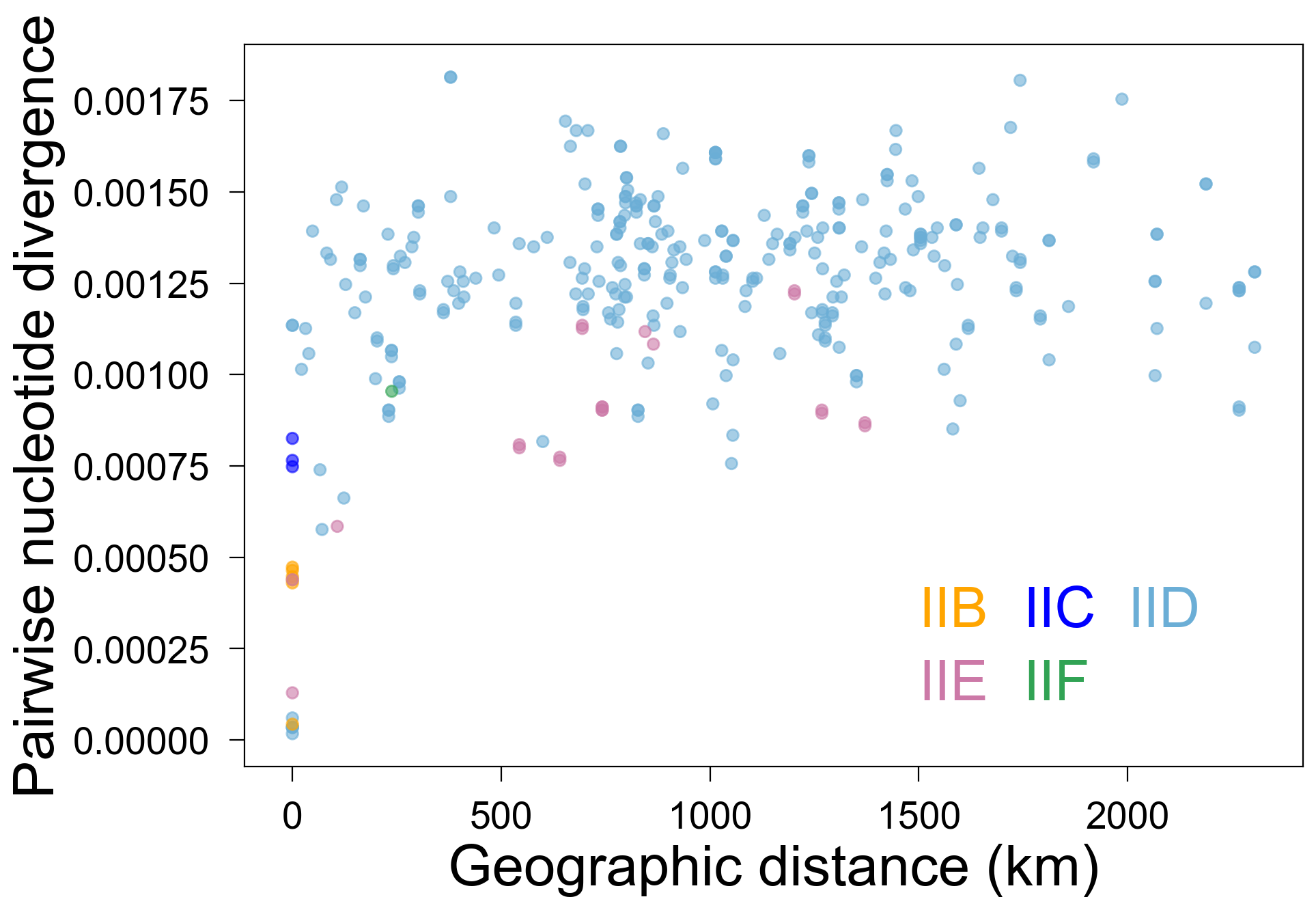

**Fig. S3. Within clonal lineages of *Ophidiomyces ophidiicola* (*Oo*), there is no evidence of recent, long-distance dispersal.** Great-circle distance (x-axis) versus pairwise nucleotide divergence (y-axis) for each pair of strains that were isolated from wild snakes and belonged to the same clonal lineage within Clade II. Each point represents a pair of strains and the colors indicate the clonal lineages. Genetic divergence was calculated using 116,322 positions in the nuclear genome that were variable among *Oo* strains from Clade II.

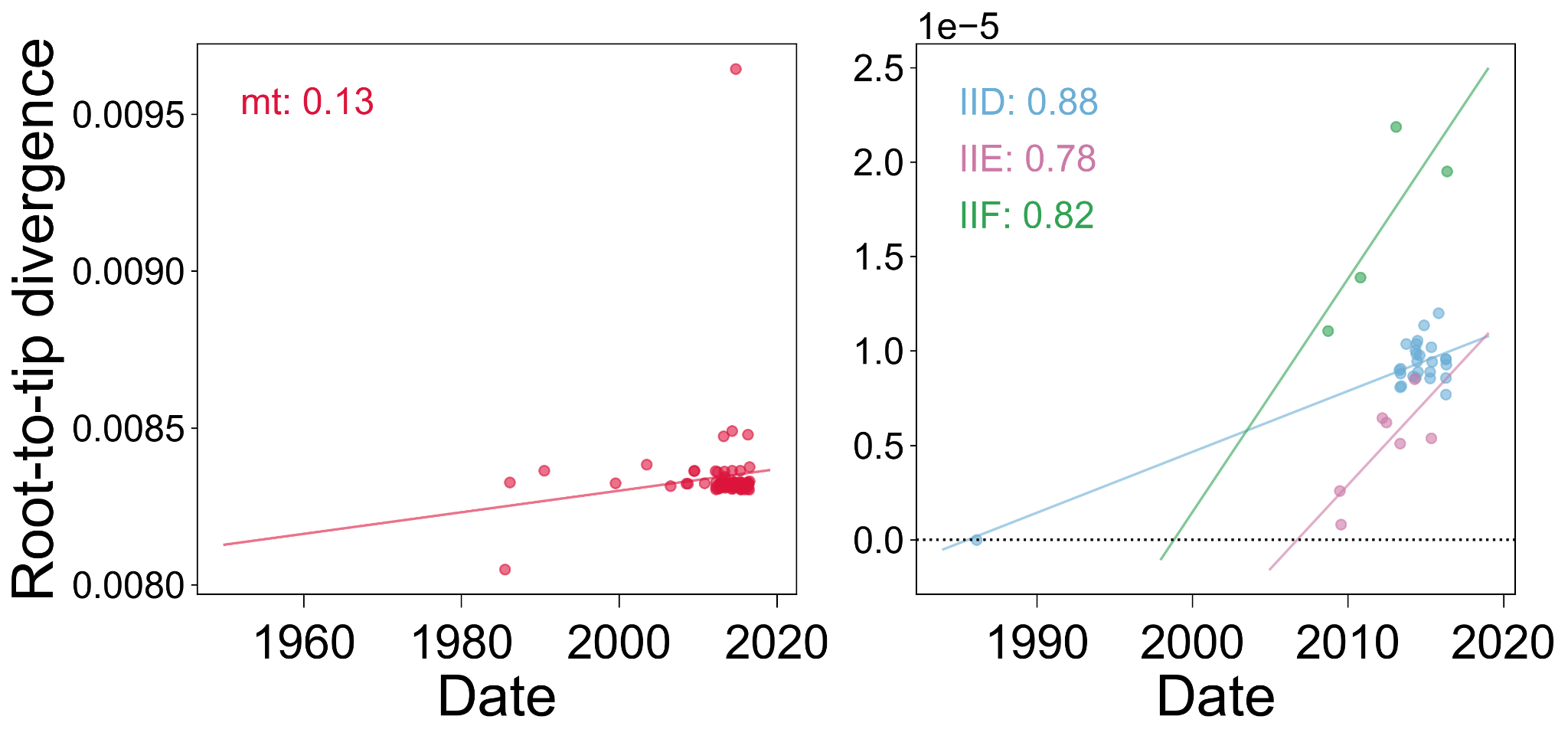

**Fig. S4.** **Root-to-tip plots demonstrate the presence of molecular clock signals within both the mitochondrial and nuclear genomes of *Ophidiomyces ophidiicola*.** Root-to-tip divergence (y-axis) was calculated using TempEst (2) v1.5.3 using maximum-likelihood phylogenies and the colored numbers inside each plot represent Pearson correlation coefficients. The mitochondrial tree (left) was rooted using the residual mean squared best-fitting root function. The nuclear trees for the Clade II clonal lineages (right) were rooted using the correlation best-fitting root function. A positive slope indicates the presence of a molecular clock signal.

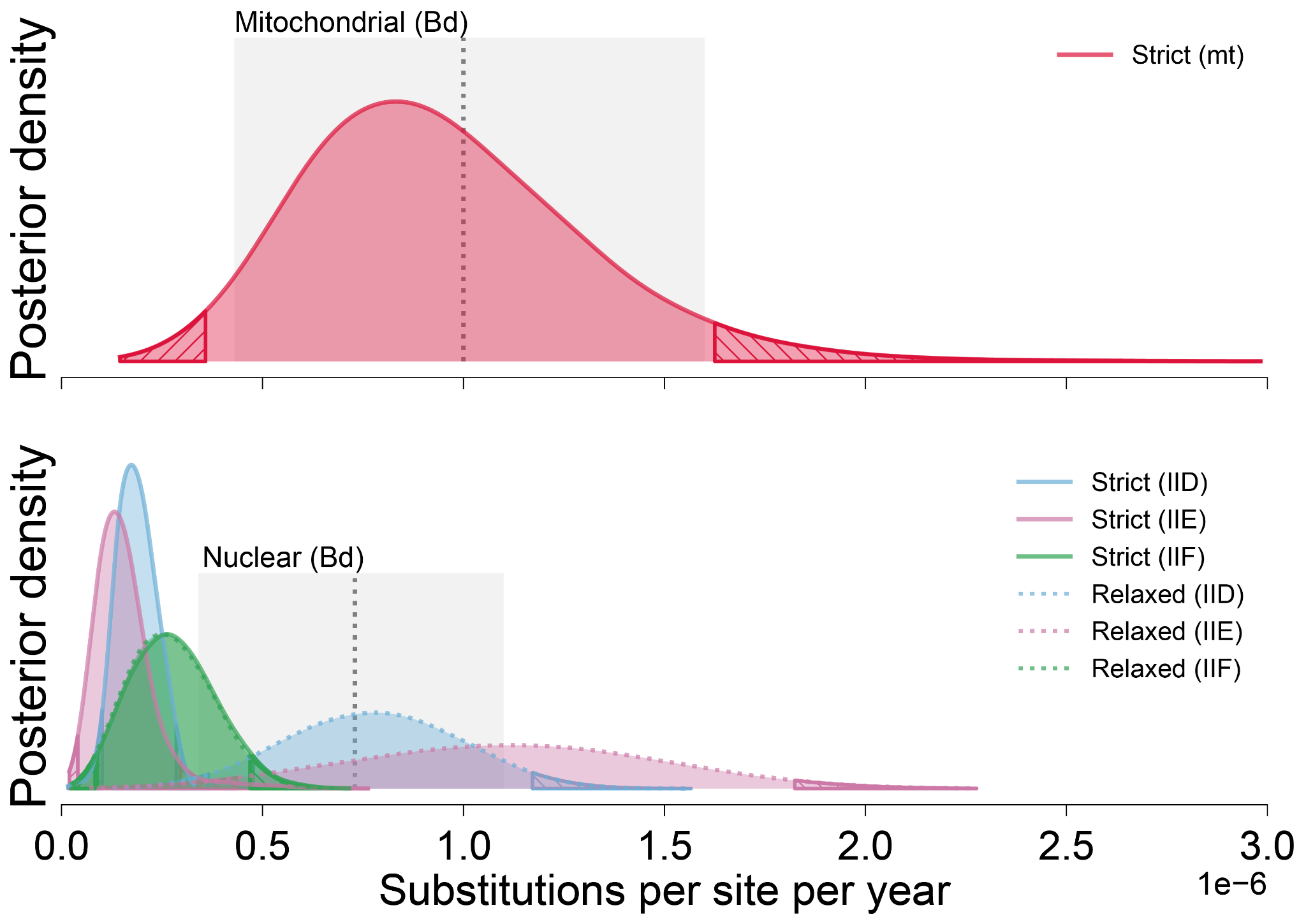
**Fig. S5. Substitution rate estimates for *Ophidiomyces ophiodiicola*** (***Oo*) are very similar to published estimates for *Batrachochytrium dendrobatidis* (*Bd*).** Colored distributions represent the posterior probability distributions from BEAST for the *Oo* substitution rate in number of substitutions per site per year. There is a single rate estimate for the mitochondrial genome based on a strict clock model (top, red) and six rate estimates for the nuclear genome, two for each of the three primary clonal lineages based on the strict (solid line) and relaxed (dotted line) clock models. Hatched regions represent the tails of each distribution that fall outside of the 95% highest posterior density. Gray boxes indicate published rate estimates for *Batrachochytrium dendrobatidis* (*Bd*) (3), with the edges of the boxes indicating the boundaries of the 95% highest posterior densities and the dashed lines representing the mean estimates.

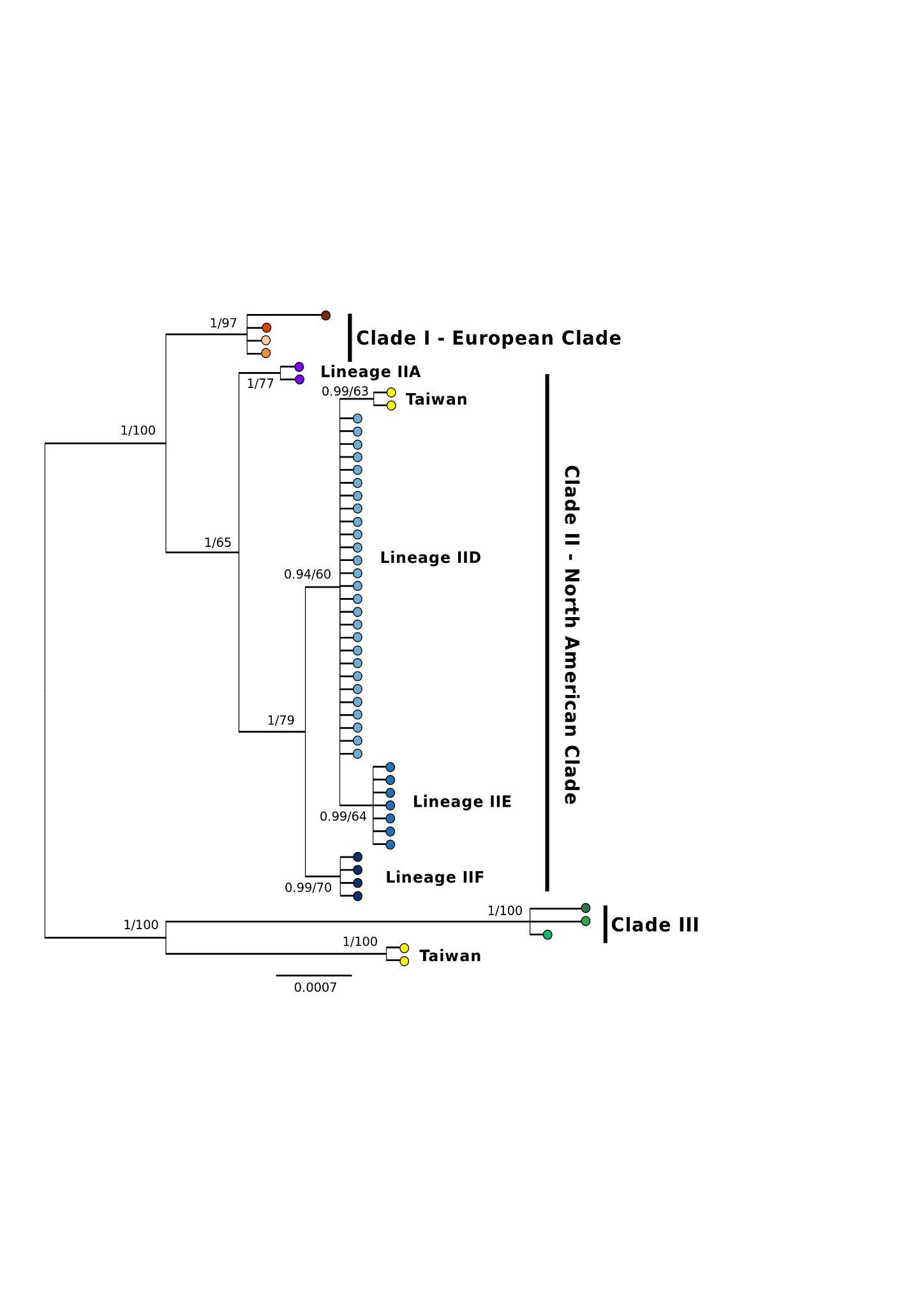

**Fig. S6.** **Phylogenetic tree of *Ophidiomyces ophidiicola* strains, including all strains from Clades I and II, nonrecombinant strains from Clade II, and strains isolated from wild snakes in Taiwan, based on three concatenated loci (internal transcribed spacer region, actin, and translation elongation factor 2ɑ).** Both Bayesian and maximum likelihood analyses produced trees with identical topologies (consensus tree from Bayesian analysis is shown). Posterior probabilities/bootstrap support values are shown at each node. Note that two strains recovered from wild snakes in Taiwan (shown in yellow) are most closely related to Clade III and two strains reside within Clade II (i.e., North American clade).

**Table S1.** Per strain summary of high-throughput whole genome sequence data and resulting *de novo* genome assemblies.

| **Strain** | **Read Length** | **Raw Reads** | **Filtered Reads** | **Average Sequencing Depth** | **De novo Contigs** | **ORFs*** |
| --- | --- | --- | --- | --- | --- | --- |
| NWHC 22687-1 | 251 | 10197907 | 10193458 | 189.93 | 127 | 6979 |
| NWHC 22747-6 | 126 | 1326261 | 1274026 | 12.79 | 805 | 6995 |
| NWHC 23906-1 | 126 | 5879326 | 5787019 | 57.20 | 224 | 6979 |
| NWHC 23913-1 | 126 | 2033427 | 2020515 | 20.50 | 511 | 7007 |
| NWHC 23942-1 | 251 | 10163217 | 10158941 | 194.44 | 95 | 7001 |
| NWHC 24042-1 | 126 | 1260472 | 1252803 | 12.94 | 840 | 7018 |
| NWHC 24216-1 | 126 | 3139116 | 3006147 | 30.11 | 335 | 6993 |
| NWHC 24266-1 | 126 | 2872471 | 2773153 | 27.87 | 360 | 7001 |
| NWHC 24266-2 | 251 | 9006524 | 9003880 | 180.99 | 102 | 6996 |
| NWHC 24266-3 | 126 | 3357160 | 3184682 | 30.15 | 426 | 6995 |
| NWHC 24266-5 | 126 | 3017486 | 3000382 | 30.64 | 360 | 7014 |
| NWHC 24266-6 | 126 | 2462441 | 2457477 | 25.27 | 436 | 6994 |
| NWHC 24281-1 | 126 | 3070566 | 2753098 | 27.95 | 995 | 7008 |
| NWHC 24392-1 | 126 | 1506101 | 1480557 | 14.57 | 672 | 7007 |
| NWHC 24393-1 | 126 | 2932825 | 2881744 | 29.99 | 326 | 7003 |
| NWHC 24395-1 | 126 | 2279108 | 2207236 | 21.82 | 399 | 7013 |
| NWHC 24411-1 | 126 | 3399511 | 2741587 | 26.93 | 357 | 7022 |
| NWHC 24414-1 | 126 | 2120595 | 2017829 | 19.32 | 449 | 7018 |
| NWHC 24415-1 | 126 | 2972904 | 2832085 | 28.67 | 408 | 7004 |
| NWHC 24564-1 | 126 | 2318899 | 2237180 | 21.80 | 538 | 7011 |
| NWHC 24746-1 | 126 | 2443737 | 2397662 | 23.58 | 360 | 7013 |
| NWHC 24824-1 | 126 | 2189724 | 2176112 | 21.31 | 496 | 7018 |
| NWHC 24825-2 | 126 | 1569775 | 1515349 | 14.81 | 581 | 6990 |
| NWHC 24828-1 | 126 | 2515775 | 2465717 | 24.66 | 427 | 7010 |
| NWHC 24852-1 | 126 | 1272897 | 1182180 | 11.59 | 818 | 6941 |
| NWHC 24874-1 | 126 | 1956670 | 1898619 | 18.33 | 502 | 7023 |
| NWHC 24878-1 | 126 | 2049710 | 2038140 | 20.38 | 422 | 7001 |
| NWHC 24878-5 | 126 | 2445299 | 2311320 | 22.94 | 1078 | 7027 |
| NWHC 24878-6 | 126 | 3594138 | 3171832 | 32.41 | 1003 | 7016 |
| NWHC 24885-1 | 126 | 2390477 | 2381952 | 23.39 | 419 | 7014 |
| NWHC 24894-1 | 126 | 1304417 | 1292860 | 12.61 | 793 | 6973 |
| NWHC 24900-1 | 126 | 2672773 | 2374420 | 23.08 | 376 | 7002 |
| NWHC 26049-1 | 126 | 2041086 | 1901559 | 18.53 | 484 | 7016 |
| NWHC 26330-2 | 126 | 1884931 | 1856116 | 17.39 | 397 | 6901 |
| NWHC 26341-1 | 126 | 2874027 | 2766065 | 27.24 | 327 | 7001 |
| NWHC 26452-1 | 126 | 1942182 | 1925839 | 18.82 | 407 | 6974 |
| NWHC 26465-1 | 126 | 3331786 | 3065353 | 30.00 | 333 | 6998 |
| NWHC 26480-1 | 126 | 1970986 | 1942893 | 18.99 | 472 | 7003 |
| NWHC 26583-2 | 126 | 2297464 | 2249150 | 22.34 | 382 | 6997 |
| NWHC 26671-1 | 126 | 3239198 | 3071511 | 30.29 | 310 | 7001 |
| NWHC 27239-1 | 126 | 2647802 | 2610585 | 25.04 | 388 | 7006 |
| NWHC 27242-2 | 126 | 2578401 | 2432908 | 24.50 | 547 | 7010 |
| NWHC 27242-3 | 126 | 3144002 | 3110471 | 31.48 | 375 | 7014 |
| NWHC 27242-4 | 126 | 3244580 | 3064536 | 30.51 | 292 | 7004 |
| NWHC 27421-1 | 126 | 3811351 | 3730730 | 38.19 | 304 | 7010 |
| NWHC 27422-1 | 126 | 2588572 | 2340580 | 23.33 | 422 | 6989 |
| NWHC 27466-1 | 126 | 4132511 | 4027173 | 40.43 | 203 | 6984 |
| NWHC 44736-14 | 126 | 2489173 | 2471815 | 23.89 | 351 | 6975 |
| NWHC 44736-31 | 126 | 2322240 | 2222581 | 22.19 | 361 | 6983 |
| NWHC 44781-7 | 126 | 2274674 | 2090969 | 20.47 | 366 | 6999 |
| NWHC 44781-8 | 126 | 1841961 | 1815989 | 17.91 | 491 | 7007 |
| NWHC 44736-45 | 126 | 1533049 | 1502150 | 14.29 | 566 | 6996 |
| NWHC 44736-52 | 126 | 2446502 | 2428252 | 24.09 | 427 | 7014 |
| NWHC 44736-65 | 126 | 1673536 | 1641537 | 16.40 | 575 | 6993 |
| NWHC 44736-75 | 126 | 1975184 | 1959985 | 19.21 | 555 | 7031 |
| NWHC 44736-81 | 126 | 1213947 | 1197527 | 11.41 | 740 | 6968 |
| NWHC 44736-83 | 126 | 2436944 | 2400837 | 24.35 | 449 | 6998 |
| NWHC 44736-86 | 126 | 2444143 | 2335495 | 23.00 | 415 | 7007 |
| NWHC 44736-87 | 126 | 2504741 | 2419170 | 24.44 | 472 | 6983 |
| NWHC 44736-88 | 126 | 2147702 | 2104036 | 20.46 | 474 | 7018 |
| NWHC 44736-89 | 126 | 2361118 | 1735994 | 16.79 | 603 | 7025 |
| NWHC 44736-90 | 126 | 2416227 | 2313226 | 23.54 | 469 | 6979 |
| NWHC 44736-93 | 126 | 3173916 | 2966920 | 28.87 | 337 | 6998 |
| NWHC 44736-94 | 126 | 2837972 | 2643284 | 26.64 | 300 | 6983 |
| NWHC 44736-95 | 126 | 1972550 | 1928576 | 18.49 | 500 | 7007 |
| NWHC 44781-25 | 126 | 1397882 | 1391262 | 13.57 | 607 | 6954 |
| NWHC 44781-26 | 126 | 2061643 | 1947185 | 18.92 | 444 | 7003 |
| NWHC 45692-2 | 251 | 9945096 | 9928858 | 191.24 | 91 | 7026 |
| NWHC 45707-81 | 126 | 1996435 | 1944233 | 18.13 | 468 | 7004 |
| NWHC 45707-82 | 126 | 1459090 | 975027 | 9.27 | 741 | 6928 |
| NWHC 45707-83 | 126 | 2636473 | 2590225 | 25.02 | 313 | 7024 |
| CBS 102663 | 126 | 1812625 | 1797218 | 17.71 | 535 | 7003 |
| CBS 122913 | 251 | 10803124 | 10793583 | 202.37 | 115 | 7007 |
| NWHC 44781-4 | 126 | 2067984 | 1948555 | 19.22 | 1352 | 6995 |
| UAMH 10296 | 251 | 9115922 | 9101483 | 198.36 | 115 | 7096 |
| UAMH 10768 | 126 | 2672744 | 2654959 | 26.79 | 324 | 6990 |
| UAMH 10949 | 126 | 2360294 | 2192868 | 21.33 | 357 | 7002 |
| UAMH 11295 | 126 | 2957143 | 2885367 | 28.20 | 265 | 6994 |
| UAMH 6218 | 126 | 2009924 | 1995802 | 19.79 | 428 | 7009 |
| UAMH 6642 | 126 | 2367804 | 2361918 | 24.05 | 380 | 6975 |
| UAMH 6688 | 126 | 3214858 | 2799322 | 32.68 | 434 | 7124 |
| UAMH 9985 | 251 | 8439071 | 8428713 | 186.24 | 133 | 7190 |
| *ORF = open reading frame | | | | | | |

**Table S2.** Metadata for strains of *Ophidiomyces ophidiicola* used in this study (see separate file).

**Table S3.** Results of PAUP* analysis for various subsets of the *Ophidiomyces ophidiicola* strains sequenced in this study.

| **Clade** | **Genome type** | **Consistency index (CI)**** | **Homoplasy index (HI)**** | **# strains included** |
| --- | --- | --- | --- | --- |
| I, II and II | Mitochondrial | 0.94 | 0.06 | 82 |
| I and II | Nuclear | 0.31 | 0.69 | 79 |
| I and IIA, IID-F | Nuclear | 0.86 | 0.14 | 44 |
| I | Nuclear | 0.99 | 0.01 | 4 |
| II | Nuclear | 0.25 | 0.72 | 75 |
| IIA, IID-F | Nuclear | 0.93 | 0.07 | 40 |
| IIA | Nuclear | N/A^ | N/A^ | 2 |
| IIB | Nuclear | 0.93 | 0.07 | 4 |
| IIC | Nuclear | N/A^ | N/A^ | 3 |
| IID | Nuclear | 1.00 | 0.00 | 27 |
| IIE | Nuclear | 0.98 | 0.02 | 7 |
| IIF | Nuclear | 1.00 | 0.00 | 4 |
| **Uninformative characters were excluded in the calculation of these metrics. | | | | |
| ^At least 4 taxa are required for unrooted tree search with PAUP. | | | | |

**Table S4.** Summary of BEAST model testing and substitution rate estimates (see separate file).

**Table S5.** Summaries of the posterior probability distributions from BEAST for the date estimates at which the most recent common ancestors of various *Ophidiomyces ophidiicola* clades existed.

| **Group** | **Genome type** | **Clock Model** | **Mean** | **Median** | **95% HPD*** |
| --- | --- | --- | --- | --- | --- |
| I, II and III | Mitochondrial | Strict | -9147.04 | -8202.62 | [-17863.2001, -2559.9348] |
| I and II | Mitochondrial | Strict | 109.57 | 271.37 | [-1362.466, 1266.7673] |
| I | Mitochondrial | Strict | 1770.30 | 1793.82 | [1552.5932, 1937.9538] |
| II | Mitochondrial | Strict | 913.78 | 1009.31 | [22.2346, 1573.3425] |
| IID-F | Mitochondrial | Strict | 1860.00 | 1875.36 | [1730.5746, 1958.2772] |
| IIE-F | Mitochondrial | Strict | 1951.17 | 1957.17 | [1902.0104, 1985.7623] |
| IID | Mitochondrial | Strict | 1951.49 | 1957.95 | [1901.7849, 1986.0562] |
| IID (wild only) | Mitochondrial | Strict | 1952.38 | 1958.58 | [1900.2787, 1992.5542] |
| IIE | Mitochondrial | Strict | 1980.67 | 1983.65 | [1961.8353, 1989.9997] |
| IIE (wild only) | Mitochondrial | Strict | 1986.69 | 1988.18 | [1963.8875, 2007.4021] |
| IIF | Mitochondrial | Strict | 1978.66 | 1983.96 | [1938.3633, 2006.5036] |
| IIF (wild only) | Mitochondrial | Strict | 1983.71 | 1989.11 | [1943.0048, 2011.5777] |
| IID | Nuclear | Strict | 1974.78 | 1976.70 | [1961.16, 1986.0784] |
| IID (wild only) | Nuclear | Strict | 1988.92 | 1990.72 | [1975.4912, 2000.7101] |
| IIE | Nuclear | Strict | 1985.41 | 1988.75 | [1959.4326, 2006.4996] |
| IIE (wild only) | Nuclear | Strict | 1985.41 | 1988.75 | [1959.4326, 2006.4996] |
| IIF | Nuclear | Strict | 1985.72 | 1989.27 | [1959.8288, 2001.2037] |
| IIF (wild only) | Nuclear | Strict | 2001.44 | 2003.23 | [1988.9764, 2009.129] |
| IID | Nuclear | Relaxed | 1975.98 | 1980.02 | [1953.0078, 1986.0837] |
| IID (wild only) | Nuclear | Relaxed | 2001.97 | 2004.16 | [1986.7355, 2011.7783] |
| IIE | Nuclear | Relaxed | 2006.98 | 2007.96 | [2001.6489, 2009.4677] |
| IIE (wild only) | Nuclear | Relaxed | 2006.98 | 2007.96 | [2001.6489, 2009.4677] |
| IIF | Nuclear | Relaxed | 1985.05 | 1988.94 | [1959.2872, 2001.8847] |
| IIF (wild only) | Nuclear | Relaxed | 2000.90 | 2002.95 | [1987.2301, 2009.1854] |
| * HPD = highest posterior density | | | | | |
| "wild only" indicates estimates that only include *Ophidiomyces ophidiicola* strains isolated from wild snakes, while other estimates may also include strains from captive snakes. | | | | | |

SUPPORTING TEXT

**Mating type locus PCR assays**

*Results*

All nine European isolates (NWHC 45692-12, NWHC 45707-84, NWHC 46078-5, NWHC 46078-6, NWHC 46078-7, NWHC 46078-8, NWHC 46078-9, NWHC 46078-10, NWHC 46078-11) screened with the primers were positive for only the *MAT1-2* locus. Amplicons were sequenced in both directions and were identical to the sequences of *MAT1-2* generated through whole genome sequencing of the other European isolates.

*Methods*

The mating type loci present in nine European isolates of *Ophidiomyces ophidiicola* (*Oo*) that were not included in the whole genome sequencing analysis were determined using newly designed PCR assays. Sequences of the *MAT1-1* and *MAT1-2* loci from whole genome sequence data of *Oo* were aligned to identify conserved regions. The program Primer3 (4-5) was then used to design two pairs of primers. The first primer pair (Oo MAT1-1 F: 5’ - GAAAGTTAAATCGGGCTTG - 3’; Oo MAT1-1 R: 5’ - TGGATGAATAGCGGTAGG - 3’) targeted a 574 nucleotide portion of the *MAT1-1* locus. The second primer pair (Oo MAT1-2 F: 5’ - CAATAGGTTGGTGCTGGT - 3’; Oo MAT1-2 R: 5’ - GTCCCAGTCGTTGCTTTC - 3’) targeted a 1,013 nucleotide portion of the *MAT1-2* locus.

The primer pairs were validated by screening 12 strains of *Oo* for which mating type was previously determined (five strains containing *MAT1-1*, seven strains containing *MAT1-2*) based on whole genome sequence data. Each reaction consisted of 23.75 μL water, 10 μL GoTaq® Flexi Buffer (Promega Corporation, Madison, Wisconsin), 5 μL 2.5 mM each dNTPs, 3 μL 25 mM MgCl₂, 2.5 μL 20 μM forward primer, 2.5 μL 20 μM reverse primer, 0.25 GoTaq® DNA polymerase, and 3 μL extracted DNA template. Cycling conditions for the PCR were as follows: 95°C for 5 min; 45 cycles of 94°C for 45 sec, 52°C for 45 sec, 72°C for 1 min; 72°C for 10 min. All strains predetermined to contain the *MAT1-1* or *MAT1-2* yielded an amplicon with the correct primer set and of the expected size.

**Phylogenetic analysis including Asian isolates of *Ophidiomyces ophidiicola***

*Results*

The four Asian isolates originated from two snakes (two isolates per individual snake). In both cases, the two isolates obtained from the same snake had identical sequences for the three loci examined. However, two unique genotypes were identified. Consensus trees resulting from the Bayesian and maximum likelihood analyses had identical topologies and good support for most of the clades and clonal lineages. One of these Asian strains (strain 2605) possessed 13 unique single nucleotide polymorphisms (SNPs) compared to the strains we sequenced. This genotype grouped with, yet was distinct from, Clade III from our analysis. The second Asian strain (strain 2664) resided within Clade II. This strain grouped with lineage IID but had three unique SNPs separating it from other isolates within IID (Fig. S6).

*Methods*

*Oo* was recently isolated from two wild snakes in Taiwan (6). However, DNA sequence data were only available for three loci (internal transcribed spacer region [ITS], actin [*ACT*], and translation elongation factor 2ɑ [*TEF*]) from those isolates; thus, those isolates could not be included in analyses using whole genome sequence data. To determine how the Asian isolates of *Oo* compared to the strains we examined, we performed a separate phylogenetic analysis by concatenating ITS, *ACT*, and *TEF* sequence data. In addition to the Asian isolates, we included all of our strains from Clades I and III and non-recombinant strains from Clade II. Sequence data for *ACT* and *TEF* were obtained from the *de novo* genome assemblies of each strain. The ITS is part of the rRNA gene complex which occurs as tandem repeats on the fungal genome and is often excluded from genome assemblies. Thus, we used ITS sequence data that was already in GenBank for a given strain or new sequence data was generated as described in previously (7). Newly generated sequences for the ITS were deposited in GenBank (see Table S2).

Sequence data for each locus were aligned using MUSCLE (8). All gaps were deleted from the alignments and data for the three loci were concatenated. For phylogenetic analyses, the loci were partitioned with a Kimura two-parameter model applied to the ITS and Hasegawa-Kishino-Yano (HKY) model applied to *ACT* and *TEF* data. A maximum likelihood analysis was performed in PAUP* v4.0a169 with 1,000 bootstrap iterations using a heuristic search technique and the sub-tree pruning and regrafting rearrangement operation. A Bayesian analysis was performed on the same dataset using MrBayes v3.2.7a through the CIPRES Science Gateway (9). For the Bayesian analysis, we conducted two runs with four chains and 5,000,000 generations, with a sampling frequency of 1,000 generations.

DISCLAIMER

Any use of trade, firm, or product names is for descriptive purposes only and does not imply endorsement by the U.S. Government.

SUPPORTING REFERENCES

1.	T. C. Bruen, H. Philippe, D. Bryant, A simple and robust statistical test for detecting the presence of recombination. *Genetics* **172**, 2665–2681 (2006).

2.	A. Rambaut, T. T. Lam, L. Max Carvalho, O. G. Pybus, Exploring the temporal structure of heterochronous sequences using TempEst (formerly Path-O-Gen). *Virus Evol* **2**, vew007 (2016).

3.	S. J. O’Hanlon, *et al.*, Recent Asian origin of chytrid fungi causing global amphibian declines. *Science* **360**, 621–627 (2018).

4.	T. Koressaar, M. Remm, Enhancements and modifications or primer design program Primer 3. *Bioinformatics* **23**, 1289-1291 (2007).

5.	A. Untergasser, *et al.*, Primer3 - new capabilities and interfaces. *Nucleic Acids Res* **40**, e115 (2012).

6.	P.-L. Sun, *et al.*, Infection with *Nannizziopsis guarroi* and *Ophidiomyces ophiodiicola* in reptiles in Taiwan. *Transbound. Emerg. Dis.* (2021). https:/doi.org/[10.1111/tbed.14049](http://dx.doi.org/10.1111/tbed.14049).

7.	J. M. Lorch, *et al.*, Snake fungal disease: an emerging threat to wild snakes. *Philos. Trans. R. Soc. Lond. B Biol. Sci.* **371** (2016).

8.	R. C. Edgar, MUSCLE: multiple sequence alignment with high accuracy and high throughput, *Nucleic Acids Res* **32**, 1792-1797 (2004).

9.	M. A. Miller, W. Pfeiffer, T. Schwartz, Creating the CIPRES Science Gateway for inference of large phylogenetic trees *in* Proceedings of the Gateway Computing Environments Workshop (GCE), 14 Nov. 2010, New Orleans, LA pp 1-8 (2010).
